# New groups of highly divergent proteins in families as old as cellular life with important biological functions in the ocean

**DOI:** 10.1101/2024.01.08.574615

**Authors:** Duncan Sussfeld, Romain Lannes, Eduardo Corel, Guillaume Bernard, Pierre Martin, Eric Bapteste, Eric Pelletier, Philippe Lopez

**Affiliations:** Institut de Systématique, Evolution, Biodiversité (ISYEB), Sorbonne Université, CNRS, Museum National d’Histoire Naturelle, EPHE, Université des Antilles, Paris, France; Génomique Métabolique, Genoscope, Institut François-Jacob, CEA, CNRS, Université d’Evry, Université Paris-Saclay, 91000 Evry, France; Research Federation for the Study of Global Ocean Systems Ecology and Evolution, FR2022/Tara Oceans GOSEE, 75016 Paris, France

**Keywords:** Microbial dark matter, Sequence similarity networks, Distant homology, Microbiome

## Abstract

**Background:** Metagenomics has considerably broadened our knowledge of microbial diversity, unravelling fascinating adaptations and characterising multiple novel major taxonomic groups, e.g. CPR bacteria, DPANN and Asgard archaea, and novel viruses. Such findings profoundly reshaped the structure of the known tree of life and emphasised the central role of investigating uncultured organisms. However, despite significant progresses, a large portion of proteins predicted from metagenomes remain today unannotated, both taxonomically and functionally, across many biomes and in particular in oceanic waters, including at relatively lenient clustering thresholds.

**Results:** Here, we used an iterative, network-based approach for remote homology detection, to probe a dataset of 40 million ORFs predicted in marine environments. We assessed the environmental diversity of 53 gene families as old as cellular life, broadly distributed across the Tree of Life. About half of them harboured clusters of environmental homologues that diverged significantly from the known diversity of published complete genomes, with representatives distributed across all the oceans. In particular, we report the detection of environmental clades with new structural variants of essential genes (SMC), divergent polymerase subunits forming deep-branching clades in the polymerase tree, and variant DNA recombinases of unknown origin in the ultra-small size fraction.

**Conclusions:** These results indicate that significant environmental diversity may yet be unravelled even in strongly conserved gene families. Protein sequence similarity network approaches, in particular, appear well-suited to highlight potential sources of biological novelty and make better sense of microbial dark matter across taxonomical scales.

## Background

Over the last decades, novel sequencing methods have allowed microbiologists to appreciate the ubiquity and abundance of uncultured organisms [1–5], and access microorganisms’ genomes beyond the isolation-cultivation dogma issued from the Koch principles [6] that underpinned microbiological studies for decades. Metagenomic studies [7] have led to an unprecedented broadening of our knowledge of microbial diversity [8], from the unravelling of microbial adaptations and interactions in numerous environments [9–12] to the characterisation of multiple novel major taxonomic groups [13–17] – most notably CPR bacteria [13, 18, 19], DPANN archaea [18, 20, 21] and Asgard archaea [22–24], profoundly reshaping the structure of the tree of life. Large groups of novel viruses [25–27] and mobile elements [28] have also been unearthed. Together, these major discoveries emphasise the central role of investigating yet uncultured organisms, believed to constitute the majority of overall microbial lineages [3, 29], in addressing many fundamental questions of biology and evolutionary biology.

Over time, as cultivation-independent sequencing efforts are carried out in an increasing range of ecosystems, discovery events of novel branches near the base of the tree of life are predicted to become less frequent [8, 17]. In accordance with this perspective, an extensive study of over 50,000 MAGs, assembled from a vast ensemble of metagenomes and including 12,556 novel candidate species-level OTUs, found no reliable evidence of novel prokaryote phylum content [30]. It may therefore seem that whatever biodiversity remains to be discovered should yield few more “major unknowns”.

However, contrasting with these observations, it still persists that across most biomes, large portions of environmental metagenomes remain taxonomically and functionally unannotated, even at relatively permissive clustering thresholds [31]. This vast pool of uncharacterised sequences remains a significant blind spot in our grasp of the extant biological diversity on Earth. Some may yet belong to genomes of unknown organisms that have so far escaped detection efforts, for instance due to accelerated evolution rates or an ancestral divergence from known organisms. Novel genes of well-characterised organisms with “open” pangenomes, divergent paralogues of known genes, and unusual mobile elements may also be expected to contribute to this “microbial dark matter” [4]. In any case, the persistence of those biological unknowns highlights the need for novel approaches complementing the current techniques to mine metagenomes for highly divergent groups.

Various network-based approaches [32], in particular, have been developed to address these concerns. Co-occurrence networks, for instance, can help assessing ecological roles of unknown taxa [33]. Sequence similarity networks, wherein pairs of primary sequences are connected according to set similarity criteria, can also be employed to compare sequences from cultured and uncultured organisms [34, 35]. In 2012, Lynch et al. used sequence similarity networks to identify several candidate new lineages from environmental 16S rRNA [36]. In 2015, Lopez et al. designed a network-based exploratory analysis to probe metagenomes for distant homologues of well-distributed gene families [37]. 86 clusters of genes broadly distributed across Domains of life were used as seeds for a two-step BLAST search inside a metagenome collection. Seed sequences were then gathered in sequence similarity networks together with their direct and indirect environmental homologues, and environmental sequences gathered in the second alignment step were more divergent from their cultured relatives than those gathered in the first round. The authors found several hundred groups of highly divergent environmental variants, some of them potentially compatible with novel major divisions of life. Consequently, (i) iterative explorations of environmental datasets may allow the retrieval of increasingly divergent variants (Fig. 1A), and (ii) network-based methods may be well-suited to handle this type of data, by integrating sequences with various levels of divergence within homologous gene families. Sequence similarity networks have also been used recently to assess how the deep-learning breakthrough in protein structure prediction may be leveraged to shed light into “functionally dark” regions of the natural protein space [38].

**Figure 1:**
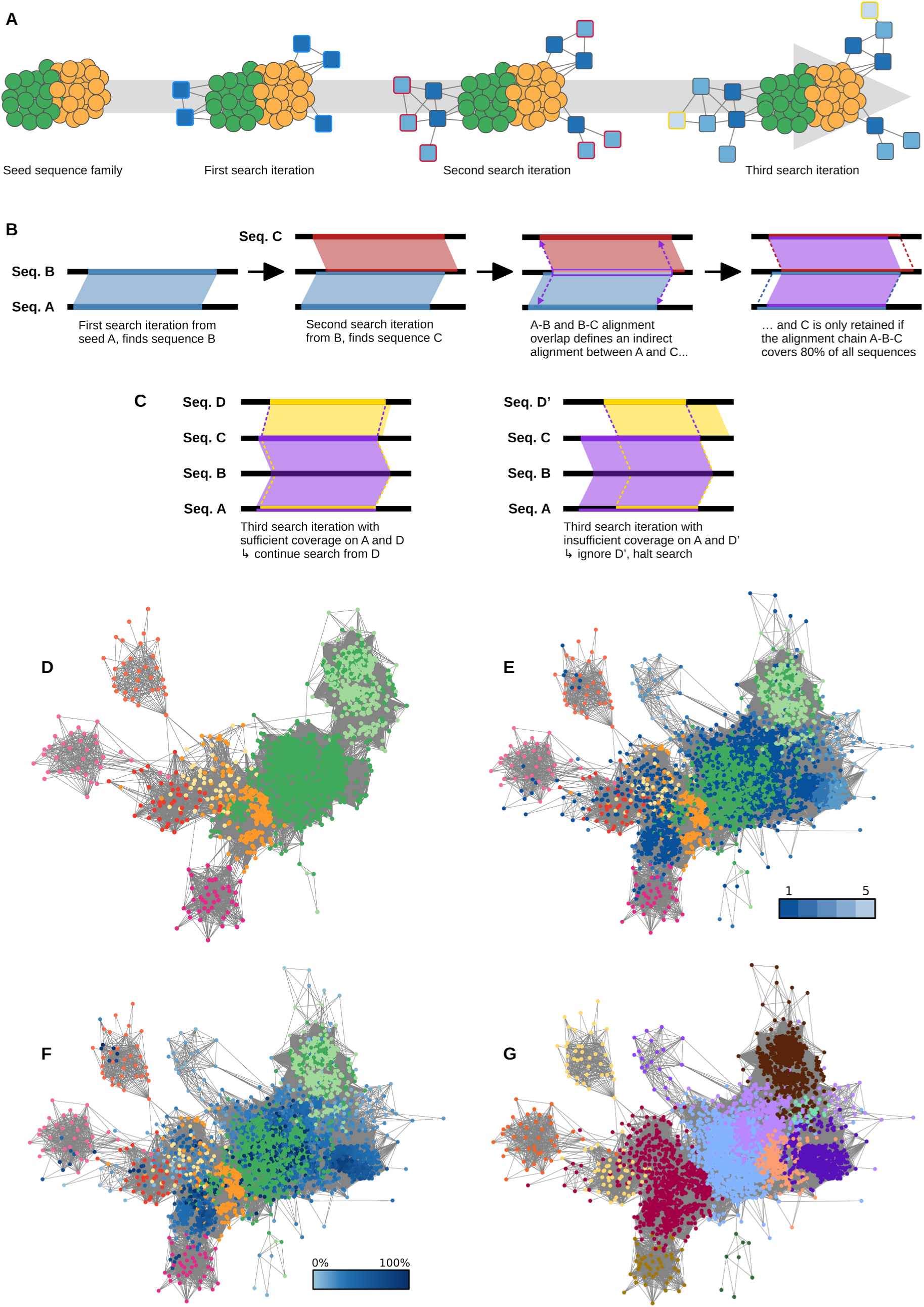
Iterative homologue search procedure. (**A**) Iterative aggregation of environmental homologues around seed sequences in a similarity network. From a set of seed sequences belonging to a given protein family (green and orange nodes), a first search iteration finds environmental homologues (dark blue nodes) for some of the seeds. A second search iteration then uses these environmental sequences as queries to find more homologues (medium blue nodes, red frame), which are themselves used as queries for a third search iteration finding further environmental homologues (light blue nodes, yellow frame). (**B**) At each iteration of the search, newly found homologues are only retained if their aligned region can be mapped back onto a seed sequence in a way that ensures >80% coverage on all sequences along the chain of aligned sequences. (**C**) **Left**: sequence D is found after three search iterations from seed A, and its alignment with sequence C can be mapped back to A in a way that preserves 80% coverage on all sequences along the “alignment chain”. Sequence D is therefore retained and will be used as query for the next iteration of the search. **Right**: sequence D’ is found after three search iterations from seed A, but its aligned region cannot be mapped back to A without breaking the 80% coverage requirement. D’ is thus not retained as a distant homologue of A in this round of search. (**D-G**) Sequence similarity networks for SMC proteins. (D) shows seed sequences only, (E-G) show seed and environmental sequences. In (D-F), nodes representing seed sequences are coloured according to their taxonomic origin (yellow: non-DPANN archaea; orange: DPANN archaea; light green: CPR bacteria; dark green: non-CPR bacteria; shades of red: four eukaryotic SMC paralogues). In (E), environmental nodes are coloured in blue, with darker shades for sequences retrieved in earlier iterations of the search, and lighter shades for sequences retrieved later. In (F), environmental nodes are coloured in blue, with darker shades for sequences with higher similarity to the known cultured diversity, and lighter shades for sequences with less similarity. In (G), all nodes are coloured according to Louvain clusters inferred in the SSN (one arbitrary colour per cluster).

In this work, we conducted an exploratory search of ocean metagenomic data to identify potential sources of novel diversity in highly conserved, near-universal gene families. Our search mined the environmental diversity of the Ocean Microbial Reference Gene Catalog (OM-RGC) dataset [39]. This extensive, non-redundant record contains sequences for over 40 million bacterial and archaeal genes, predicted from metagenomic sequencing of a large variety of marine environments across the world. At the time of initial publication, around 45% of these sequences lacked taxonomical annotation at or below the Domain level, and 43% lacked functional annotation to an eggNOG orthologous group (OG), highlighting the existence of a vast, undescribed diversity in the global oceanic microbiome, as well as the necessity of additional efforts to improve its characterisation. To perform this search, we further developed the iterative explorative strategy of environmental datasets initiated by Lopez et al. [37], by allowing distant homologue search iterations to continue indefinitely until convergence. Specifically, we focussed our search on ancestral gene families that showed particular conservation across their taxonomic distribution in the face of evolution. Retrieving highly divergent variants in such families could indeed carry an increased biological significance, given their stability in primary sequence for many reference genomes, and potentially guide future searches for novel putative taxonomical groups or biological functions involving these nearly universal gene families. We thus used a custom dataset of 53 ancient, conserved gene families with key biological functions to initiate our iterative probing of OM-RGC. We identified highly divergent variants of multiple gene families, uncovering new putative structural and sequence variants of biologically essential proteins across taxonomical scales.

## Results and Discussion

### Oceanic metagenomes harbour distant homologues of highly conserved protein families

We developed an iterative mining procedure to accumulate highly divergent environmental variants for families of genes or proteins of interest. From an initial set of nearly ten million protein sequences gathered from prokaryotic, eukaryotic, viral and plasmidic complete genomes (Table SI-1), we selected a set of 53 protein clusters, highly conserved and at least as old as cellular life. Most clusters corresponded to single protein families, though a few of them comprised proteins from two or more closely related families (we hereafter refer to those clusters as families for simplicity, and will make multiplicity cases explicit when discussing such clusters specifically). These families spanned a total of 125,774 sequences and included 12 families of ribosomal proteins (Table SI-2). On average, bacterial sequences in these families had 34.9% amino-acid identity to their closest archaeal homologue (and vice-versa), roughly illustrating the level of divergence to expect between sequences from different Domains of life.

Each selected family was used as the seed for a deep homologue-mining procedure in the OM-RGC dataset [39]. This iterative search aimed at aggregating around each seed family the diversity of its environmental homologues, including variants too divergent to produce a significant direct alignment to any seed sequence. For each family, direct oceanic homologues of seed sequences were identified in a first round of search. The OM-RGC dataset was then further queried for homologues of those homologues, and so forth until the procedure converged to find no additional environmental homologues (See Fig. 1A-E and Methods for details).

We tested the performance of our method by conducting homology searches on a simulated dataset, and found that our protocol was particularly resistant to false-positive homology calls. More specifically, we sought to evaluate (i) how reliably our iterative procedure successfully retrieved distant homologues of seed sequences, and (ii) whether this retrieval was prone to false-positive calls, where sequences would be attained from seeds that did not share a homologous origin. To that end, we generated a collection of phylogenetic trees, based on a common balanced binary tree structure, where branches along the path from the root to another given node (internal or terminal) were elongated to represent various levels of divergence. Each tree was then assigned a randomly generated amino-acid sequence, which was evolved numerically along its branches, resulting in some slow- and some fast-evolving terminal sequences. Slow-evolving sequences within the same “family” shared an average of 42.7% sequence identity. Finally, the slow-evolving subsets were each used as seeds for iterative homology searches to retrieve their own fast-evolving homologues amongst all sequences generated from all phylogenies. Across the 3402 test cases that were performed in total, we detected no instance of false-positive homology hit, i.e. homology searches only ever retrieved sequences genuinely related to the seeds. In cases where fast-evolving sequences diverged up to 2.5 times faster than their slow counterparts, the search procedure was nearly systematically able to retrieve all divergent sequences (Fig. SI-1). When the evolution rate difference was four-fold, about half of the test instances successfully retrieved all divergent homologues. Finally, above a six-fold increase, seed sequences were largely unable to retrieve any divergent sequence at all. These results on simulated data show that the procedure we developed to identify remote homologies aggregates new sequences in an efficient but conservative manner that resists spurious homology calls, although the higher complexity of real-world biological sequence data may be expected to yield aberrant results on occasion.

Our iterative metagenome mining procedure expanded the selected 53 seed families by a total of 826,717 environmental sequences from OM-RGC (Fig. SI-2). All seed families had their own set of environmental homologues, requiring an average of 7 rounds of iterative search before exhaustion. Despite metagenomic sequencing sometimes yielding shorter gene sequences than what is anticipated from genomes in culture, sequences retrieved from OM-RGC were only marginally shorter than their reference counterparts (Pearson *r*=0.96, p-value 3.5×10^-30^), further confirming that their divergence was not related to a systematic bias associated with sequence size.

OM-RGC homologues of the 53 selected seed families were then compared against proteins from the NCBI non-redundant (*nr*) database to find their closest relative amongst <u>all</u> published sequences with taxonomically resolved annotations (Fig. 1F; Supplementary Text SI-1). Only 6.7% of all retrieved environmental sequences were >90% similar to their closest characterised relative, implying that a large majority of environmental proteins cannot be accurately represented by genomes captured by current cultivation or isolation techniques. Furthermore, 20.5% of environmental variants had less than 34.9% similarity with their closest *nr* relative, i.e. they diverged more from any proteins of well-characterised organisms than bacterial and archaeal homologues diverged from one another on average in the reference dataset. Environmental homologues of ribosomal protein families had generally higher similarity to their closest characterised relative than non-ribosomal environmental sequences (one-sided Kolmogorov-Smirnov test, p-value <1.6×10^-22^; Fig. SI-3), possibly owing to their reputedly high evolutionary conservation. Still, even ribosomal protein families included very divergent oceanic variants (Fig. SI-3). Moreover, all sampled oceanic sites revealed similar proportions (but uneven absolute numbers) of divergent and highly divergent prokaryotic sequences (Fig. SI-2). Any location in the global ocean could therefore be a prolific reserve of new microbial gene variants, including temperate surface-layer habitats. Some of the retrieved environmental sequences show levels of divergence to the known diversity that are comparable with the difference between archaeal and bacterial homologues. These variants could potentially belong to uncharacterised lineages that branched away from well-known taxa long ago, although alternative hypotheses can be offered: divergent environmental homologues could, for instance, be distant paralogues of seed sequences, that evolved faster than their known counterparts due to relaxed selective pressure after duplication, and appear environmentally conserved but not described in cultured organisms.

### Highly divergent clusters of environmental variants expand the diversity of multiple universal protein families

Seed sequences and their (direct and indirect) oceanic homologues were then gathered in family-specific sequence similarity networks (SSNs). Similar sequences in these networks are expected to gather in coherent, well-connected groups, thus reflecting the structure of protein families in the network topology. Sequences within each SSN were therefore partitioned into network communities using Louvain clustering [40] (Fig. 1G). This higher-level view of network structures allows an easier assessment of the environmental diversity, including identifying potential sources of biological novelty in these protein families. In particular, clusters consisting exclusively or predominantly of environmental sequences (>90% of environmental sequences), with little similarity to published sequence records (<40% sequence identity to any non-environmental sequence in the nr database), and containing enough proteins to be unlikely the result of sequencing inaccuracies, are intuitively the most likely to correspond to genuinely novel groups of environmental homologues.

691 clusters of sequences were inferred in total across the 53 SSNs, of which we retained 80 clusters of proteins fitting the above criteria for significant novelty potential. These 80 clusters of highly divergent sequences were distributed across 25 ancient, conserved protein families. Remarkably, no cluster with such a high level of divergence was found in networks of ribosomal proteins, possibly due to a superior level of conservation or a higher coverage of their diversity in public sequence databases. Still, the fact that clusters of divergent environmental homologues were identified in nearly half of our selected protein families suggests that numerous key biological processes are carried out by a currently underestimated diversity of protein primary structures. In other words, the “functional dark matter” of proteins likely consists of both unknown functions and unknown actors of known functions [31, 41].

To assess how these groups of divergent sequences may relate to their reference counterparts, we reconstructed phylogenetic trees regrouping seed and environmental sequences from each of the 80 selected highly divergent clusters. This selection exposed an additional phylogenetic diversity in conserved protein families when environmental contributions are considered. In particular, in some families, sequences representative of certain divergent network clusters branched between or beside the main groups of archaeal and bacterial sequences. Such phylogenetic placements indicate substantial potential for novelty in the sequence space of those protein families. We detail findings of particular interest for three families in the following subsections.

#### High environmental diversity in oceanic DNA polymerase clamp loaders

One of the selected seed families in our study consisted of several AAA+ ATPases [42], mostly involved in clamp-loading systems for DNA replication. In the environmental diversity of this family, we identified large contingents of highly divergent variants across the phylogeny of the family.

In a mechanism conserved across all cellular life forms, DNA polymerases process and replicate DNA by binding onto circular clamps that encircle and slide along the template DNA strand. Sliding clamps are embedded onto DNA by a pentameric clamp-loading system, which exhibits a universally conserved structure in archaea, bacteria and eukaryotes despite differences in subunit composition [43]. All clamp loaders consist of one “large” subunit (δ in bacteria, RfcL in archaea, Rfc1 in eukaryotes) complemented by four “small” subunits: three γ and one δ’ subunits in bacteria (also respectively called DnaX and HolB), four RfcS subunits in archaea, one each of Rfc 2-5 subunits in eukaryotes. All subunits are homologous to one another within and across all three Domains of life [44–47].

Our seed family consisted of sequences for the clamp loader “small” subunits (CLSSUs) described above (i.e. bacterial DnaX and HolB, archaeal RfcS, and eukaryotic Rfc 2-5), as well as sequences for the bacterial replication-associated recombination protein RarA. This protein, present in bacteria and eukaryotes but not in archaea [48], is involved in homologous recombination and DNA repair, both in the context of DNA replication and outside [49]. The RarA protein sequence is highly conserved and also substantially homologous to DnaX, and as such was grouped alongside it in the construction of our seed families.

The iterative retrieval of environmental homologues for this protein family resulted in a nearly five-fold increase of its sequence content (Table SI-2). In particular, the resulting SSN harboured 10 new clusters of highly divergent environmental homologues (Fig. SI-4). Owing to their high divergence in primary sequence, not all clusters translated to perfectly monophyletic groups in the phylogeny we produced (Fig. 2), though they still generally maintained some level of coherence. Amongst the ten environmental clusters, one had its representative sequences branch within reference archaeal and eukaryotic Rfc sequences (cluster 26), and another translated to a new clade within reference HolB/DnaX bacterial sequences (cluster 23). Additionally, one environmental cluster branched next to bacterial RarA sequences (cluster 27), and its sequences were annotated as belonging to the B subunit of the Holliday junction resolving complex RuvABC, already shown to cluster near clamp-loading proteins in sequence networks [50]. Finally, sequences from seven divergent clusters resulted in groups <u>outside</u> the bacterial and archaeal/eukaryotic seed sequence clans [51] in the phylogeny (clusters 2, 14, 15, 16, 19, 24, 25). Eggnog annotations for these sequences mapped them predominantly to HolB (COG0470), though it should be noted that one particular cluster contained 96% of functionally unassigned sequences (cluster 24).

**Figure 2:**
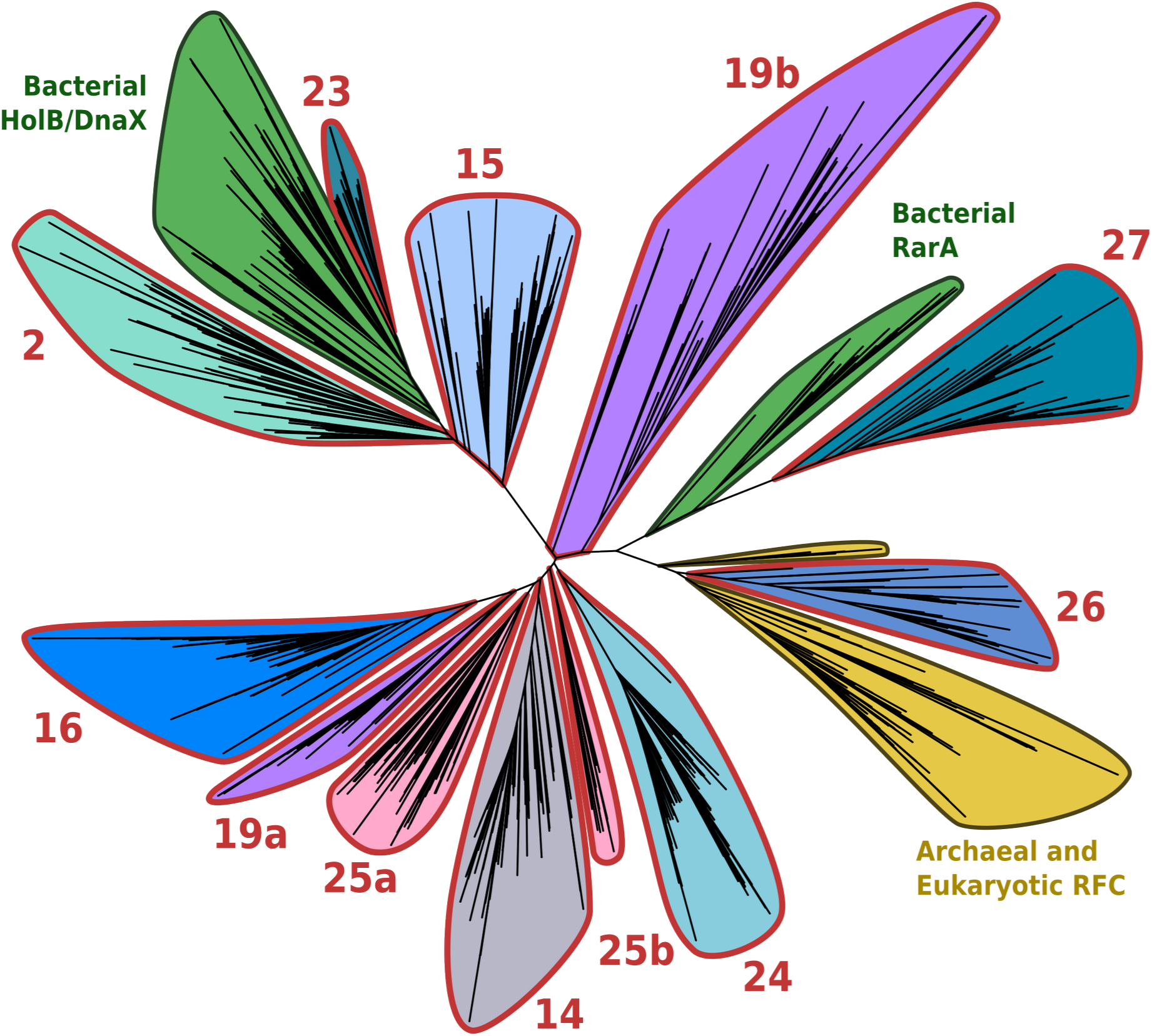
Alignment-free phylogeny of the DNA clamp loader subunits: HolB/DnaX/RarA/RFC sequences and environmental homologues from significantly divergent clusters. Seed sequences are coloured according to the Domain of life of their host organism (green: Bacteria, yellow: Archaea and Eukaryotes). Groups of environmental sequences are coloured according to the network cluster they belong to in the family SSN, and outlined in red. Numerical cluster labels are inherited from Fig. SI-4 and shared with Fig. 3. Note: environmental network clusters 19 and 25 are both split into two groups in this phylogenetic tree.

Protein structures were predicted for representatives of seed and divergent environmental CLSSUs using ColabFold [52, 53], and gathered in a dendrogram depicting their similarities (Fig. 3). Most seed proteins used for this comparison showed similar structures, although HolB, DnaX, RarA and archaeal/eukaryotic Rfc still formed distinct groups in the structure dendrogram. Structures inferred from environmental variants followed a pattern similar to the sequence phylogeny, with representatives from clusters 2, 15 and 23 branching near HolB references, and most other clusters translating to structures sitting outside of the main reference groups. In other words, the environmental HolB variants that we identified on the basis of primary sequence divergence also exhibited a divergence in 3D structure consistent with their phylogenetic placements.

**Figure 3:**
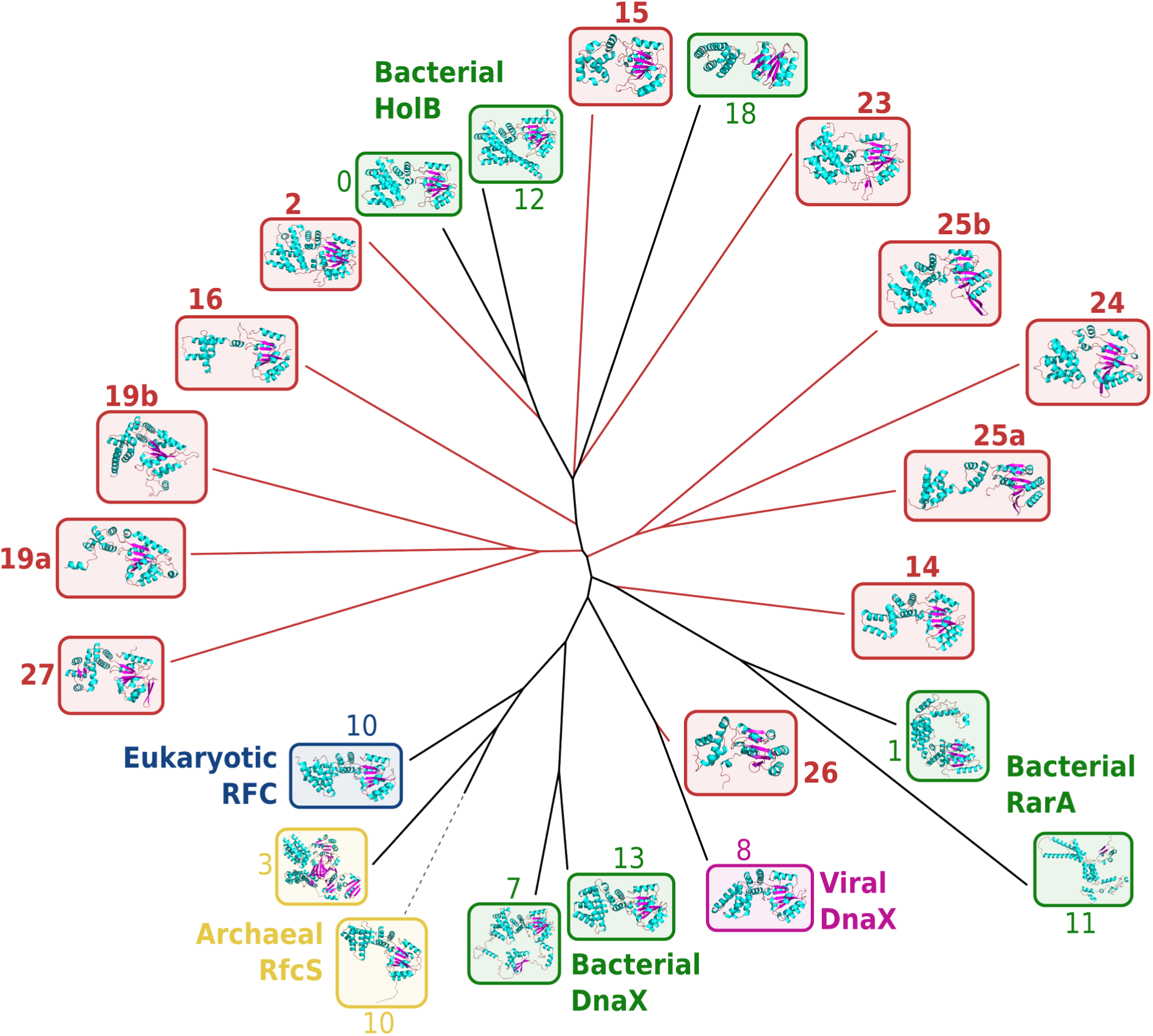
Dendrogram of tertiary structures of DNA clamp loader subunits: HolB/DnaX/RarA/RFC sequences and environmental homologues from significantly divergent clusters. Protein structures were inferred with AlphaFold and compared (all against all) using Foldseek. Leaves and structures are boxed according to the Domain of life of their host organism (green: Bacteria, yellow: Archaea, magenta: Viruses). Environmental leaves and structures are boxed in red, with numerical labels corresponding to the SSN cluster they belong to, in accordance with Fig. 2 and Fig. SI-4.

Oceanic homologues of CLSSUs therefore diverge from their known counterparts in primary sequence, and exhibit tertiary structures comparable, but not identical, to canonical CLSSU structures. Such structural differences and sequence-based phylogenetic placements for these highly divergent environmental CLSSU homologues could reflect the existence of undetected divergent paralogues in these gene families, which would raise interesting questions about their possible contribution in such a conserved subprocess of DNA replication. These could also hypothetically be indicative of some unknown microbial lineage(s), though much more conclusive data would be required before firmly asserting this. In any case, these results hint at a diversity of uncultivated marine organisms replicating DNA using various unusual proteic machineries, possibly resulting in unusual replication mechanisms operating in the ocean.

#### Novel abundant clade of SMC proteins with unusual structure in Actinobacteria

Another remarkable seed family consisted of SMC (structural maintenance of chromosomes) proteins, and we identified a small but abundant group of environmental SMC variants with strikingly singular structures within Actinobacteria.

SMC proteins are present in all Domains of life and act (as part of the SMC complex) as regulators of high-order chromosome organisation [54]. Eukaryotic genomes encode six paralogous SMC proteins (SMC1-6), due to a sequence of duplications around the time of the last eukaryotic common ancestor. Indeed, a single copy of the *smc* gene is present in nearly all archaea and bacteria, with a few exceptions. In some γ-proteobacteria a different proteic complex, MukBEF, is responsible for these functions instead [55]. Bacteria from various phyla can also harbour another complex, MksBEF, alongside their SMC or MukBEF machinery [56]. MksBEF is believed to be evolutionarily related to MukBEF, and both are structurally analogous to the SMC complex, but primary sequence comparisons have ruled this structural similarity as convergent rather than due to distant homology [54]. SMC complexes are also notably absent from Crenoarchaeota, resulting in distinctive chromosomal dynamics and cell cycle logics [57, 58].

A typical SMC protein consists of five domains: an N-terminal domain containing a Walker A motif; a first helical chain of roughly 300 amino-acids; a central “hinge” domain; a second α-helix of comparable length to the first; a C-terminal domain containing a Walker B motif [59, 60]. This linear structure self-folds by linking the N- and C-terminal motifs into an ATPase “head”, with the two α-helix domains forming an antiparallel coiled-coil between this head and the hinge domain. This hinge then serves as a dimerisation site for a second SMC monomer, with accessory proteins binding to the ATPase heads to complete the ring-shaped SMC complex [54]. The hinge region of the SMC complex subsequently plays the essential role of mediating DNA binding, and allows the loading of SMC rings onto chromosomes [61, 62].

From seed sequences in this family, we retrieved a rather limited amount of environmental homologues (0.97 environmental homologue per seed sequence in this family, compared to a median value of 2.6 across all families, see Table SI-2), but one small cluster of distant environmental homologues was still identified (cluster 9 in Fig. SI-5). In the phylogeny produced from seed SMC sequences and oceanic variants from this cluster (Fig. 4), environmental sequences formed a monophyletic clade branching close to the base of seed actinobacterial sequences. These divergent environmental sequences were functionally annotated as SMC proteins (COG1196), and were strikingly abundant in the sequencing data, nearly seven times more so than other OM-RGC SMC homologues. Moreover, this novel oceanic clade harbours SMC-related proteins that are critically different in structure from canonical SMC proteins (Fig. 5A; average TM-score between two proteins in the divergent cluster: 0.828; average TM-score between a protein in the divergent cluster and a reference SMC protein: 0.440). Namely, these oceanic variants lack the hinge domain which is normally essential to SMC assembly and function (Fig. 5B). As such, they may be considered more similar to bacterial SbcC and archaeal and eukaryotic Rad50 proteins, thought to be distant evolutionary relatives of SMC [54]. Indeed, proteins from this ancestral family also consist of an SMC-like head and an antiparallel coiled-coil with no hinge domain, dimerising instead through a zink-hook structure induced by a CXXC motif [63]. However, FoldSeek structural comparisons clearly discriminate between reference Rad50/SbcC proteins on one side, and SMC proteins (reference or divergent OM-RGC variants) on the other (Fig. SI-6). The zink-hook CXXC motif conserved in Rad50/SbcC is also absent from our environmental cluster sequences, confirming them as divergent variants within the SMC diversity rather than beside it.

**Figure 4:**
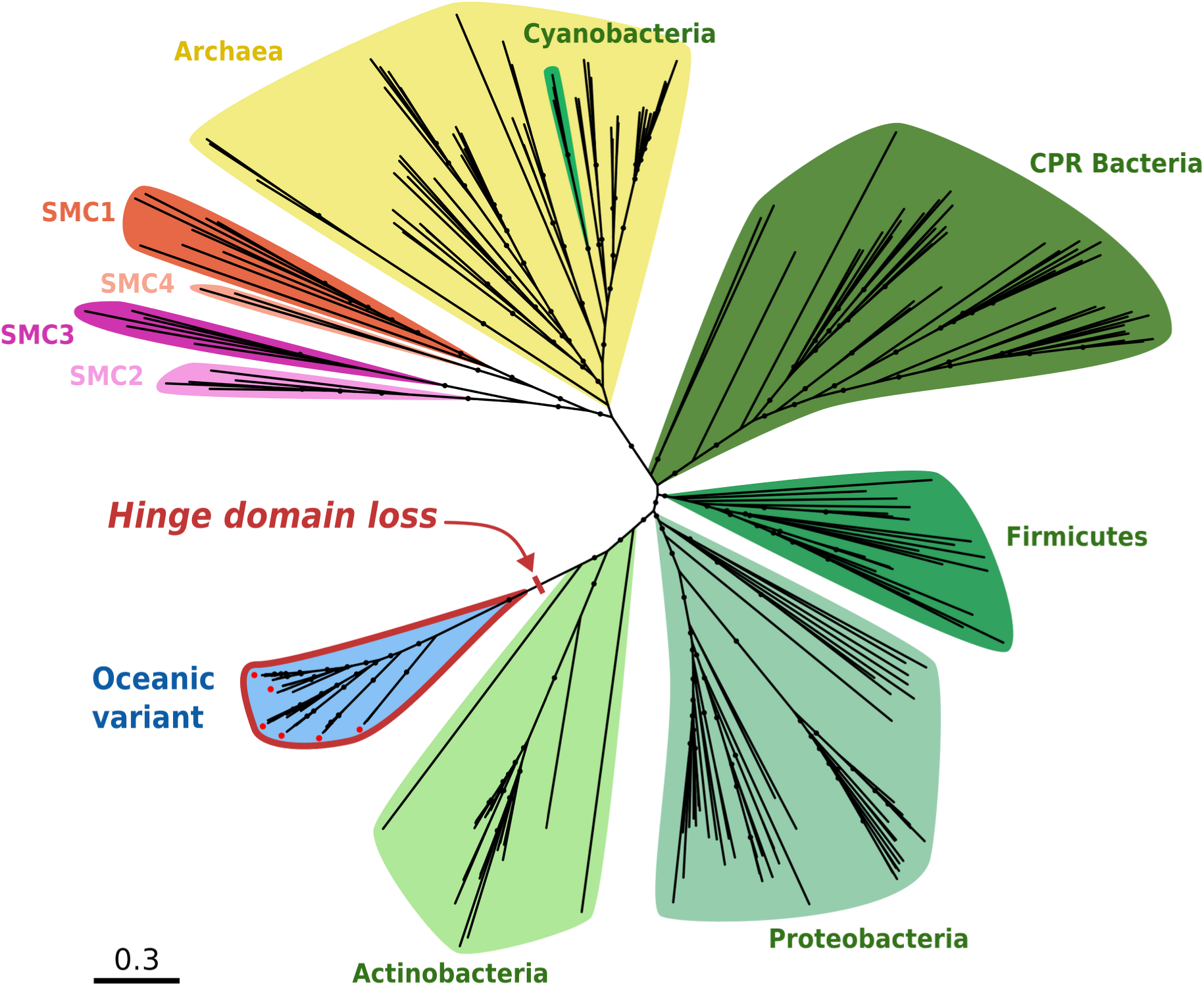
Maximum likelihood phylogenetic tree of SMC sequences and environmental homologues from significantly divergent clusters. Seed sequences are coloured according to the Domain of life of their host organism (green tones: Bacteria, yellow: Archaea, orange and purple tones: Eukaryotes). Environmental sequences are coloured in blue and outlined in red. Red dots indicate environmental sequences for which 3D structures were inferred. Black dots indicate branches with >85% bootstrap support.

**Figure 5:**
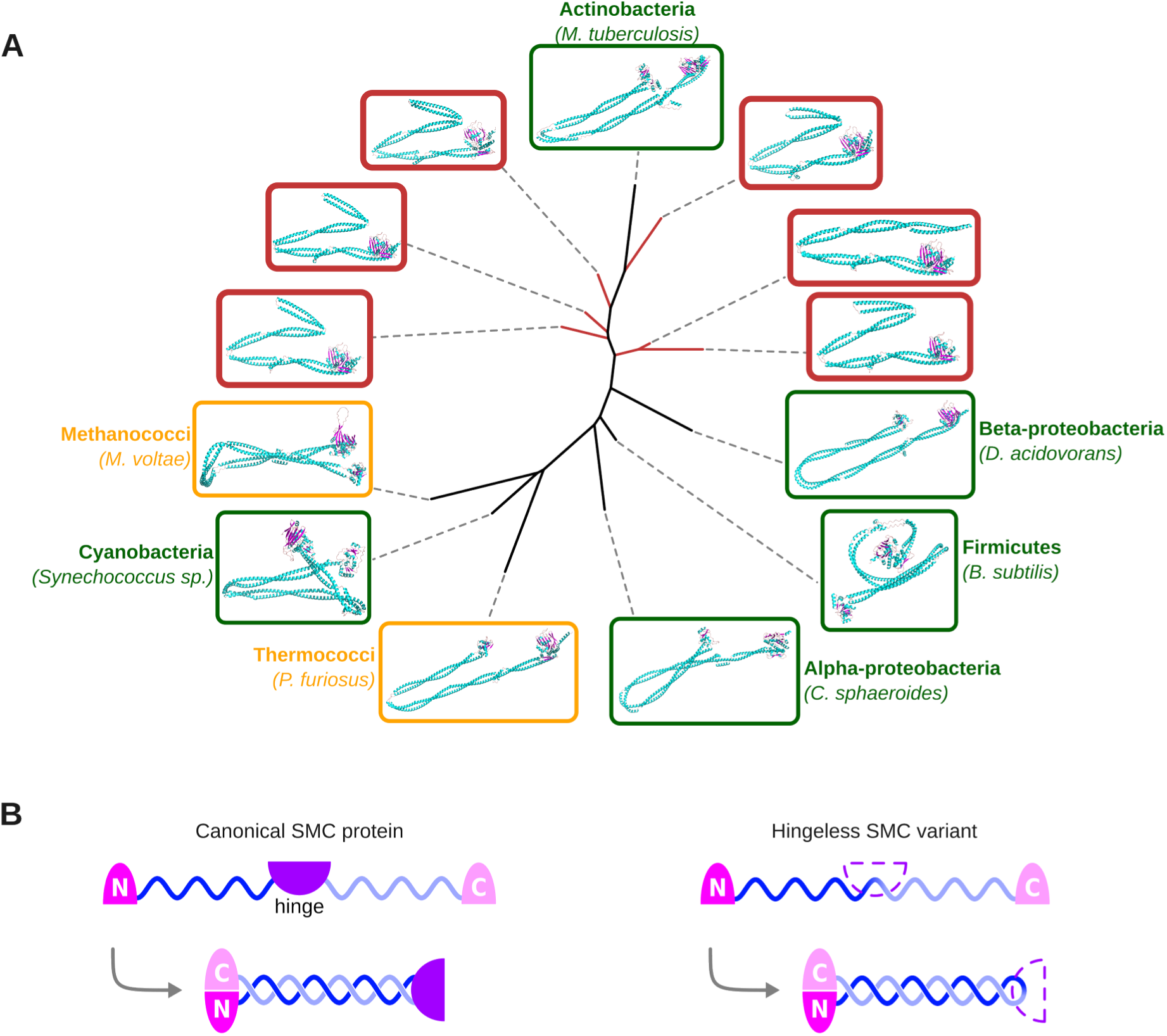
Environmental SMC homologues with divergent tertiary structure. (**A**) Dendrogram of tertiary structures of SMC sequences and selected environmental homologues from significantly divergent clusters. Protein structures were inferred with AlphaFold and compared (all against all) using Foldseek. Leaves and structures are boxed according to the Domain of life of their host organism (green: Bacteria, yellow: Archaea). Environmental leaves and structures are highlighted in red. (**B**) Schematic structure of SMC monomers. Left: canonical SMC protein with N- and C-terminal ATP-binding motifs, linked to a central hinge domain by two coiled-coil regions. This linear structure folds (grey arrow) by joining the two terminal motifs into an ATPase domain, forming a helical coiled-coil with the arm regions between the ATPase and hinge domains. Right: “hinge-less” environmental SMC homologue lacking a hinge domain. The folded protein still features the ATPase domain at one end of the coiled-coil helix, without the hinge at the opposite end.

Several evolutionary scenarios could explain this new bacterial cluster of “hinge-less” SMC. Firstly, it could be indicative of some paralogue of SMC existing in Actinobacteria. This would then be, to the best of our knowledge, the first description of SMC duplication in prokaryotes [64]. Alternatively, this divergent cluster could indicate the existence of an unknown lineage, supposedly branching within Actinobacteria, where the SMC hinge domain would have been lost. In any case, the substantial divergence of these environmental sequences to any gene published from a well-characterised organism, together with the loss of the essential hinge domain and their remarkably high abundance in the sampling data, suggests that we identified a new kind of biology within the SMC family. By the absence of their expected interaction site with DNA, one would speculate that these hinge-less SMC-related proteins must either perform a different function than known SMC or bind DNA through different mechanisms. The broad distribution of hinge-less SMC variants across the oceans, their monophyly and their relative abundance in the ocean microbiome suggest that they play an important, underappreciated function in this oceanic clade.

#### Divergent recombinases from potentially novel groups in sub-micrometre size fractions

In a third family, consisting of RecA/RadA DNA recombinases [65], we identified other possible sources of novel diversity, including within ultra-small cell size fractions.

During the course of DNA replication, accidental double-strand breaks (DSBs) in the DNA molecule can have detrimental effects on genome stability and cell viability [66]. Recombinase proteins in the RecA/RadA family are central to homologous recombinational repair, a key replicative stress-reduction pathway that can correct DSBs as well as other types of DNA damage. This family contains the extensively studied bacterial recombinase RecA (also present in eukaryotic organelles) as well as its archaeal and eukaryotic homologues, respectively RadA and Rad51 [65, 67–69].

Identifying distant environmental homologues of this seed family increased its total size five-fold (Table SI-2). Amongst this added diversity, four clusters of environmental sequences were retained as highly divergent, totalling 1700 sequences. A phylogenetic tree was produced from seed sequences as well as representative sequences for these divergent environmental clusters (Fig. 6). In this phylogeny, sequences from a first cluster branched near the root of archaeal seed sequences, and was functionally categorised as RadA (COG1066) in accordance with this placement (cluster 20 in Fig. SI-7). A second cluster of divergent environmental sequences (cluster 12) branched within the environmental ultra-small cluster in Bacteria. Interestingly, this cluster was predominantly annotated as ArlH (COG2874), an archaeal protein involved in the biogenesis of the archaellum, a cellular motility structure analogous to bacterial flagella [70]. Structure and sequence similarities between ArlH and bacterial RecA have previously been described [71] but, to the best of our knowledge, no evolutionary hypothesis has yet been put forth to explain this surprising homology. Finally, one cluster of distant environmental RecA homologues (COG0468) branched within bacterial sequences (cluster 5), and a final cluster, also annotated as RecA, saw its representative sequences sit between the archaeal and bacterial references (cluster 19). Interestingly, both of these clusters were composed of >50% of sequences from the “ultra-small” size fraction of cells with diameters <0.2 µm. Such cellular sizes are akin to those of CPR bacteria and DPANN archaea [72]; however, seed sequences from these ultra-small superphyla branch clearly within the clans of their respective Domains of life. Additionally, environmental sequences from these ultra-small clusters bore no remarkable similarity to viral sequences recorded in the NCBI Virus sequence database (accessed in February 2023) and just 11 of them (out of 1700) matched to a single oceanic virus from the GVMAG database [73].

**Figure 6:**
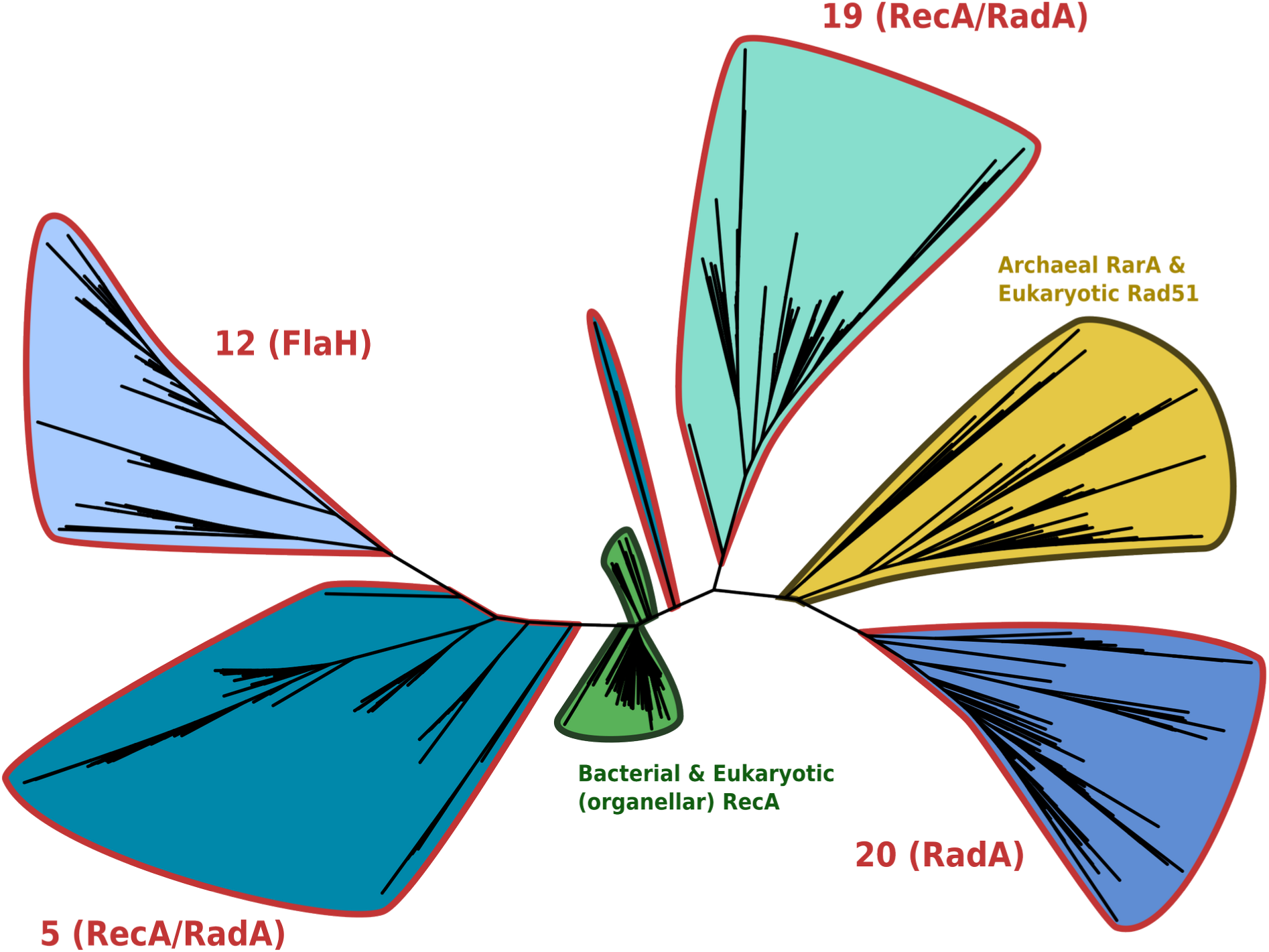
Alignment-free phylogeny of RecA/RadA sequences and environmental homologues from significantly divergent clusters. Seed sequences are coloured according to the Domain of life of their host organism (green: Bacteria and eukaryotic organelles, yellow: Archaea and eukaryotic nuclei). Groups of environmental sequences are coloured according to the network cluster they belong to in the family SSN, and outlined in red. Numerical cluster labels are inherited from Fig. SI-7.

The origin of these divergent RecA proteins therefore remains open: they could, for instance, belong to unknown bacteriophages or mobile elements populating the global ocean. They might also result from a duplication and divergence of the *recA* gene in CPR bacteria, although they should in that case be expected to appear in published genomes for members of this lineage. They may yet genuinely belong to new uncharacterised, deep-branching cellular lineages of sub-micrometre cell size, though significantly more evidence would again be required to support this hypothesis. Nevertheless, our finding of new very deep-branching groups related to RecA is consistent with the description of new basal groups of metagenomic RecA sequences formerly proposed [74], and highlights the ultra-small size fraction as a notable source of novelty in this essential protein family. Uncovering divergent forms of RadA in metagenomes is also exciting, because even some forms of RadA previously described as inactivated have been demonstrated to be functionally relevant for their host cells, and putatively attached to an alternative mechanism of replication initiation or in the regulation of origin recognition [75]. Moreover, sequence divergence, typically in the non conserved region of intein-containing RadA, may be functional, as it may affect the temperature-induced splicing of the intein of RadA, a phenotype that has been described in *Thermococcus sibericus* [76].

## Conclusion

The prevalence of biological unknowns in environmental metagenomes remains, to this day, vast; vast indeed to the extent that “known unknowns” and “unknown unknowns” constitute a relevant distinction to address genes, organisms, processes and interactions at play in the uncultured microbial world. With our network-based, multi-marker iterative approach, we sought to understand the structure of environmental genetic variation for a range of ancient, conserved gene families with functions essential to cellular life. We found that environmental variants for those gene families could exist in marine microbiomes with considerable divergence to the known diversity. Moreover, these highly divergent sequences organised in (sometimes vast) cohesive groups of homology, supposedly harboured in (sometimes vast) groups of related genomes, as illustrated by the oceanic variants of DNA polymerase clamp loaders, hinge-less SMCs, and deep-branching divergent RecA/RadA variants from the ultra-small size fraction.

A common issue surrounding metagenomic data is to know whether predicted genes and proteins actually exist in the sampled environment or result from aberrations in the assembly process. To avoid this pitfall, we purposefully limited our analyses to larger clusters of (similar but non-identical) sequences, from the already non-redundant OM-RGC dataset. Furthermore, the nature of our retrieval process imposes at least 80% of the length of any retrieved sequence to map back to at least 80% of a seed sequence (Fig. 1B-C). As such, recombined proteins mixing sequence fragments from several protein families are unlikely to be matched to our “canonical” seed families if exogenous regions cover more than 20% of their length. Lastly, the benchmarks we performed on simulated protein families show that sequences unrelated to the search seeds are seldom retrieved by erroneous homology calls. For these reasons, we believe that the groups of oceanic variants we discussed correspond to genuine environmental homologues of reference sequences, rather than assembly artifacts, protein recombinants, or non-homologous proteins from unrelated families.

Still, various competing scenarios of evolution and diversification could explain highly divergent homologues such as those we detected. We list some of them here, understanding that a single “one-size-fits-all” explanation to all the divergent groups we identified is highly unlikely.

A first hypothesis could be that an environmental cluster represents deep paralogues resulting from an ancestral duplication in the gene family. Though not impossible, this hypothesis does require an explanation as to why these paralogues do not appear more broadly in the wide range of public genomes currently available, save for some unlikely event of widespread parallel gene loss across the tree of life. Alternatively, these divergent sequences could have been spawned by more recent gene duplications at narrower taxonomic scales, after which they would have diverged rapidly from their “original” copy. This is entirely possible, predominantly for clusters clearly branching inside the phylogenetic clade of established taxa. The divergent SMC proteins we identified within Actinobacteria are perhaps an example of this (this would then be the first description of an SMC duplication in prokaryotes), though once again it would leave unexplained why most actinobacterial genomes do not seem to carry these “hinge-less” variants. Cases like this are also interesting from a functional standpoint, as the rapid divergence in primary sequence following gene duplication raises questions of neo- or subfunctionalisation for the novel paralogue.

Divergent homologues of highly conserved, ancestral families could also stem from uncharacterised genomes bearing these variants. Marine viruses, or other mobile elements, could be carrying such variants, especially those identified in smaller organism size fractions, such as the divergent forms of recombinase A we reported. It is possible that the divergence of these homologues could then point to radical gene changes, driven by specific selective pressures associated with non-cellular organisms. Conversely, unknown cellular lineages that diverged recently (e.g. from known genera or families) could also harbour unusual gene variants. In the functions we specifically targeted, strong constraints on sequence evolution are expected, meaning that drastic changes in intracellular processes or external selective pressure may have prompted those high levels of sequence divergence over short evolutionary timeframes. Lastly, the levels of divergence observed from some environmental groups could be compatible with novel major taxonomic groups that diverged from the established diversity some hundreds of millions, or even billions of years ago. This last hypothesis would, of course, require a lot more evidence to substantiate such a claim, and full genomes with high levels of divergence across their length would have to be produced and analysed thoroughly. Still, however remote, the possibility for new basal branches in the tree of life should not be fully discarded in the absence of conclusive evidence favouring other hypotheses.

All in all, the detection of divergent variants in key protein families, that have likely existed since cellular life began, supports the notion that major gaps remain in our knowledge of biological diversity, and that various forms of exciting new biology may be expected from unravelling this microbial world. To that end, future methodological extensions that rely less on primary sequence comparisons still appear warranted to address the whole natural diversity. The recent breakthroughs in protein structure prediction, in particular, could greatly benefit microbial dark matter analyses, as 3D structures tend to be more conserved than primary sequences during evolution. As such, the development of 3D similarity networks, connecting protein structures from cultured organisms to structures predicted from metagenomes, could offer unprecedented insights into the evolution and the functional landscape of environmental microbiomes, with possible applications to fields such as ecological, biotechnological or biomedical sciences.

## Materials & Methods

### Constitution of a conserved protein families dataset

We constituted a dataset of 9,737,821 proteins, from 4403 bacterial (including CPR), 567 archaeal (including DPANN and Asgard), 120 eukaryotic, 18,020 viral and 1586 plasmidic genomes, acquired from public NCBI databases [77] (Table SI-1). The sequence similarity network (SSN) of this protein collection was reconstructed by an all-against-all DIAMOND blastp alignment [78] (version 2.0.9, thresholds: E-value ≤10^-5^, sequence identity ≥30%, mutual coverage ≥80%). This SSN contained 891,459 protein clusters (connected components). The assortative mixing between Domains of life within each cluster was computed using the Python package networkx [79] (version 2.8.8). We retained 53 protein clusters meeting thresholds of (i) Domain assortativity ≥0.65 and (ii) 150 or more sequences from both archaea and bacteria. These 53 protein families comprised a total of 125,774 sequences.

### Iterative retrieval of environmental homologues

40,154,822 gene sequences from the Ocean Microbial Reference Gene Catalog (OM-RGC v1) [39] were collected, alongside corresponding sampling metadata and eggNOG [80] annotations, and translated into amino-acid sequences. An iterative search for environmental homologues in the OM-RGC dataset was conducted for the selected 53 protein families independently (building upon [37]). For each family, seed sequences were aligned against the OM-RGC protein sequences with DIAMOND (thresholds: E-value ≤10^-5^, sequence identity ≥30%, mutual coverage ≥80%). Environmental sequences retrieved were used as a base for a new round of DIAMOND alignment (identical parameters) against OM-RGC. This procedure was iterated, each round using as queries the environmental sequences retrieved in the previous round, until no additional sequence was found (Fig. 1A). At each step, the aligned regions of matched sequences were checked to project back to a region covering at least 80% of a seed sequence, to maintain the plausibility of distant homology between indirectly linked sequences (Fig. 1B-C). Sequences not meeting this criterion were discarded before the next search iteration. 826,717 sequences in the OM-RGC dataset were assigned to the selected protein families in this way (Table SI-2).

### Precision and accuracy of our iterative retrieval protocol on simulated protein families

From a balanced binary tree with 64 leaves, we generated a collection of toy phylogenies. For each non-root node in the starting tree, new trees were created by elongating branches between the root and this node, by a factor of 1 (“null” case), 1.5, 2, 2.5, 3, 3.5, 4, 6 or 8, yielding 126 non-root nodes × 9 possible elongation factors = 1134 (non-unique) tree instances. Random sequences of 300 amino-acids were then generated and numerically evolved along the branches of these trees using pyvolve [81] (version 1.0.3, LG model). Doing three replicates per tree instance, we thus simulated a total of 3402 artificial protein families with 64 members each.

In each tree we generated, branches were only elongated from the root to one target node, and therefore only on one side of the root, leading to leaf nodes on that side being further away from the root than the leaves on the opposite side. Sequences simulated along those trees could therefore be classified as slow- or fast-evolving depending on their side in the tree. 3402 iterative homology searches (same parameters as for real-world data) were thus conducted, each time using the slow-evolving sequences from one simulated family to find their fast-evolving homologues within the entire set of generated sequences. The precision (percentage of true positive homology calls amongst all retrieved sequences) and recall (percentage of fast-evolving homologues successfully retrieved) of the search protocol were determined from these results, for each possible factor of divergence, and each possible depth in the tree this divergence spanned (from 1, stopping at a node directly under the root, to 6, all the way to a leaf node).

### Comparison of retrieved environmental sequences to cultured diversity

Environmental sequences retrieved for each of the 53 selected seed families were compared to published sequences from taxonomically-resolved organisms in the NCBI *nr* database (downloaded in March 2020) via a DIAMOND alignment search (E-value ≤10^-5^). Similarity values between environmental sequences and their closest published relative were calculated as the product of the amino-acid identity in the aligned region times the alignment coverage on the shortest sequence.

### Sequence similarity network reconstruction and analysis

SSNs were computed for each environmentally expanded protein family by conducting all-against-all DIAMOND blastp alignments of seed and environmental sequences (E-value ≤10 ^-5^, sequence identity ≥30%, mutual coverage ≥80%). We then inferred, using Louvain clustering (implemented in networkx, v2.8.8) [40], node communities in those networks, i.e. groups of sequences tightly connected by homology links. This clustering defined 691 communities across the 53 families in our dataset. We further selected clusters containing at least 30 sequences, of which at least 90% were from the environmental dataset, and with environmental sequences averaging 40% identity or less with their closest published counterpart. 80 such clusters were identified across 25 families.

SSNs were rendered using Cytoscape (version 3.9.1) [82]. However larger networks, typically with millions of edges, made visualisations intractable. Synthetic “meta-networks” of those SSNs were created instead (Fig. SI-4, SI-5, SI-7). Rather than showing interconnections between all sequences, these represented connections between sequence clusters (as defined above): each Louvain cluster inferred in an SSN was condensed to a single “meta-node”, and two meta-nodes were linked by a “meta-edge” if the corresponding clusters were adjacent in the SSN. Meta-edges were also given a numeric weight representing the proportion of edges between clusters, relative to the total possible number of edges if the clusters had been fully connected together.

### Phylogenetic analysis of divergent clusters

Sequences from divergent clusters were gathered in phylogenetic trees along with seed sequences. We used CD-HIT (90% identity threshold, version 4.8.1) [83] to dereplicate sequences from each of the 80 selected clusters, as well as seed sequences from each of the 25 corresponding families. Up to 100 sequences per environmental cluster and 200 seeds per family were selected as representatives. We then first computed cluster-specific maximum likelihood phylogenies. Sequences from each divergent cluster were aligned with corresponding seed sequences using Mafft (version 7.520, 1000 iterative refinement cycles) [84]. These alignments were then trimmed using trimAl (version 1.4.1) [85], and phylogenies were produced using IQ-TREE (version 1.6.12, 1000 bootstrap replicates) [86–88]. Next, we inferred family-wide alignment-free phylogenies, grouping together (representatives of) seed sequences and all divergent clusters from each family [89, 90]. *k*-mer-based distance matrices were computed between all representative sequences of a family using jD2Stat (version 1.0, *k*=7) [91], and used to infer Neighbour-Joining trees with RapidNJ (version 2.3.2) [92]. All trees were rendered and annotated in iTOL (version 6.9) [93].

### Inferrence and comparison of protein tertiary structures

3D structures were inferred for a selection of representative sequences in the SSNs of SMC proteins and DNA clamp-loading subunits.

For clamp loaders, one sequence was selected as representative for each cluster in the SSN. Divergent environmental clusters were represented by the environmental sequence with the highest degree (number of edges in the SSN) to other environmental sequences within the cluster; other clusters were represented by the reference sequence with the highest degree to other references in the cluster. For SMC proteins, which have a significantly longer primary sequence (around 1200 amino-acids), we sought to reduce the number of structures to infer *de novo.* Six sequences from the divergent cluster of environmental SMC variants were chosen arbitrarily (all had maximal degree, because the cluster was fully connected), and public AlphaFold structures [53, 94] were acquired from UniProt [95] to represent reference SMC sequences (UniProtKB accessions: P9WGF2, Q5N0D2, A3PMS2, A9BZW2, P51834, Q69GZ5, Q8TZY2) and their Rad50/SbcC homologues (UniProtKB accessions: A0A7I7YPX7, A5GLL1, O68032, A0A210VWK9, A0A640H0H1, P62134, P58301).

Structures were inferred for selected clamp loaders and environmental SMC sequences using ColabFold (v1.5.2, default parameters) [52]. Then, reference and environmental clamp loader structures were compared using FoldSeek (version 7-04e0ec8, all-against-all, easy-search mode, no pre-filter, alignment by TM-Align) [96]. Inferred environmental SMC structures were compared with UniProt reference SMC structures following the same protocol. For both protein families, these comparisons were used to construct dendrograms with RapidNJ [92], taking as distance metric between two structures the average local distance difference test (lDDT) score of the corresponding bidirectional structural alignment. Dendrograms were plotted in iTOL [93] and annotated with 3D models of the protein structures rendered by PyMOL (version 2.5.5).

## Declarations

### Ethics approval and consent to participate

Not applicable.

### Consent for publication

Not applicable.

### Availability of data and materials

The OM-RGC dataset analysed in this study is available from this webpage: http://ocean-microbiome.embl.de/companion.html. The dataset of conserved protein families constructed for this analysis is available from this Figshare project: https://doi.org/10.6084/m9.figshare.24893910.v1. Source code for the iterative retrieval of environmental homologues can be found on the following repository: https://github.com/TeamAIRE/SHIFT.

### Authors’ contributions

DS conducted the formal analyses, with RL contributing to the creation of the seed dataset. DS, RL, EC, GB and PM contributed to methods design and software implementations. EB, EP and PL designed and supervised the study. DS wrote the original draft, with the help of EB, EP and PL to edit and finalise the manuscript. All authors read and approved the final manuscript.

## Acknowledgements

The authors are grateful to Charles Bernard for his insights on designing the iterative search procedure.

## References

1. Staley JT, Konopka A. Measurement of in situ activities of nonphotosynthetic microorganisms in aquatic and terrestrial habitats. Annu Rev Microbiol. 1985;39:321–46.

2. Amann RI, Ludwig W, Schleifer KH. Phylogenetic identification and in situ detection of individual microbial cells without cultivation. Microbiol Rev. 1995;59:143–69.

3. Whitman WB, Coleman DC, Wiebe WJ. Prokaryotes: The unseen majority. Proc Natl Acad Sci. 1998;95:6578–83.

4. Marcy Y, Ouverney C, Bik EM, Lösekann T, Ivanova N, Martin HG, et al. Dissecting biological “dark matter” with single-cell genetic analysis of rare and uncultivated TM7 microbes from the human mouth. Proc Natl Acad Sci U S A. 2007;104:11889–94.

5. Alain K, Querellou J. Cultivating the uncultured: limits, advances and future challenges. Extremophiles. 2009;13:583–94.

6. Koch R. Untersuchungen uber bakterien V. Die aetiologie der milzbrand-krankheit, begrunder auf die entwicklungegeschichte Bacillus anthracis. Beitrage Zur Biol Pflanz. 1877;2:277–310.

7. Handelsman J, Rondon MR, Brady SF, Clardy J, Goodman RM. Molecular biological access to the chemistry of unknown soil microbes: a new frontier for natural products. Chem Biol. 1998;5:R245–9.

8. Castelle CJ, Banfield JF. Major New Microbial Groups Expand Diversity and Alter our Understanding of the Tree of Life. Cell. 2018;172:1181–97.

9. Delmont TO, Robe P, Cecillon S, Clark IM, Constancias F, Simonet P, et al. Accessing the Soil Metagenome for Studies of Microbial Diversity. Appl Environ Microbiol. 2011;77:1315– 24.

10. Ventosa A, de la Haba RR, Sánchez-Porro C, Papke RT. Microbial diversity of hypersaline environments: a metagenomic approach. Curr Opin Microbiol. 2015;25:80–7.

11. Behzad H, Gojobori T, Mineta K. Challenges and Opportunities of Airborne Metagenomics. Genome Biol Evol. 2015;7:1216–26.

12. Sunagawa S, Acinas SG, Bork P, Bowler C, Eveillard D, Gorsky G, et al. Tara Oceans: towards global ocean ecosystems biology. Nat Rev Microbiol. 2020;18:428–45.

13. Hugenholtz P, Pitulle C, Hershberger KL, Pace NR. Novel division level bacterial diversity in a Yellowstone hot spring. J Bacteriol. 1998;180:366–76.

14. Chouari R, Le Paslier D, Dauga C, Daegelen P, Weissenbach J, Sghir A. Novel major bacterial candidate division within a municipal anaerobic sludge digester. Appl Environ Microbiol. 2005;71:2145–53.

15. Pelletier E, Kreimeyer A, Bocs S, Rouy Z, Gyapay G, Chouari R, et al. “Candidatus Cloacamonas Acidaminovorans”: Genome Sequence Reconstruction Provides a First Glimpse of a New Bacterial Division. J Bacteriol. 2008;190:2572–9.

16. Hug LA, Baker BJ, Anantharaman K, Brown CT, Probst AJ, Castelle CJ, et al. A new view of the tree of life. Nat Microbiol. 2016;1:1–6.

17. Parks DH, Rinke C, Chuvochina M, Chaumeil P-A, Woodcroft BJ, Evans PN, et al. Recovery of nearly 8,000 metagenome-assembled genomes substantially expands the tree of life. Nat Microbiol. 2017. 10.1038/s41564-017-0012-7.

18. Rinke C, Schwientek P, Sczyrba A, Ivanova NN, Anderson IJ, Cheng J-F, et al. Insights into the phylogeny and coding potential of microbial dark matter. Nature. 2013;499:431–7.

19. Brown CT, Hug LA, Thomas BC, Sharon I, Castelle CJ, Singh A, et al. Unusual biology across a group comprising more than 15% of domain Bacteria. Nature. 2015;523:208–11.

20. Huber H, Hohn MJ, Rachel R, Fuchs T, Wimmer VC, Stetter KO. A new phylum of Archaea represented by a nanosized hyperthermophilic symbiont. Nature. 2002;417:63–7.

21. Baker BJ, Comolli LR, Dick GJ, Hauser LJ, Hyatt D, Dill BD, et al. Enigmatic, ultrasmall, uncultivated Archaea. Proc Natl Acad Sci. 2010;107:8806–11.

22. Spang A, Saw JH, Jørgensen SL, Zaremba-Niedzwiedzka K, Martijn J, Lind AE, et al. Complex archaea that bridge the gap between prokaryotes and eukaryotes. Nature. 2015;521:173–9.

23. Zaremba-Niedzwiedzka K, Caceres EF, Saw JH, Bäckström D, Juzokaite L, Vancaester E, et al. Asgard archaea illuminate the origin of eukaryotic cellular complexity. Nature. 2017;541:353–8.

24. Imachi H, Nobu MK, Nakahara N, Morono Y, Ogawara M, Takaki Y, et al. Isolation of an archaeon at the prokaryote–eukaryote interface. Nature. 2020;577:519–25.

25. Paez-Espino D, Eloe-Fadrosh EA, Pavlopoulos GA, Thomas AD, Huntemann M, Mikhailova N, et al. Uncovering Earth’s virome. Nature. 2016;536:425–30.

26. Zhou Y, Zhou L, Yan S, Chen L, Krupovic M, Wang Y. Diverse viruses of marine archaea discovered using metagenomics. Environ Microbiol. 2023;25:367–82.

27. Gaïa M, Meng L, Pelletier E, Forterre P, Vanni C, Fernandez-Guerra A, et al. Mirusviruses link herpesviruses to giant viruses. Nature. 2023;616:783–9.

28. Al-Shayeb B, Schoelmerich MC, West-Roberts J, Valentin-Alvarado LE, Sachdeva R, Mullen S, et al. Borgs are giant genetic elements with potential to expand metabolic capacity. Nature. 2022;610:731–6.

29. Lloyd KG, Steen AD, Ladau J, Yin J, Crosby L. Phylogenetically Novel Uncultured Microbial Cells Dominate Earth Microbiomes. mSystems. 2018;3.

30. Nayfach S, Roux S, Seshadri R, Udwary D, Varghese N, Schulz F, et al. A genomic catalog of Earth’s microbiomes. Nat Biotechnol. 2020;39:499–509.

31. Bernard G, Pathmanathan JS, Lannes R, Lopez P, Bapteste E. Microbial Dark Matter Investigations: How Microbial Studies Transform Biological Knowledge and Empirically Sketch a Logic of Scientific Discovery. Genome Biol Evol. 2018;10:707–15.

32. Liu Z, Ma A, Mathé E, Merling M, Ma Q, Liu B. Network analyses in microbiome based on high-throughput multi-omics data. Brief Bioinform. 2021;22:1639–55.

33. Zamkovaya T, Foster JS, de Crécy-Lagard V, Conesa A. A network approach to elucidate and prioritize microbial dark matter in microbial communities. ISME J. 2021;15:228–44.

34. Forster D, Bittner L, Karkar S, Dunthorn M, Romac S, Audic S, et al. Testing ecological theories with sequence similarity networks: marine ciliates exhibit similar geographic dispersal patterns as multicellular organisms. BMC Biol. 2015;13:16.

35. Arroyo AS, Iannes R, Bapteste E, Ruiz-Trillo I. Gene Similarity Networks Unveil a Potential Novel Unicellular Group Closely Related to Animals from the Tara Oceans Expedition. Genome Biol Evol. 2020;12:1664–78.

36. Lynch MDJ, Bartram AK, Neufeld JD. Targeted recovery of novel phylogenetic diversity from next-generation sequence data. ISME J. 2012;6:2067–77.

37. Lopez P, Halary S, Bapteste E. Highly divergent ancient gene families in metagenomic samples are compatible with additional divisions of life. Biol Direct. 2015;10:64.

38. Durairaj J, Waterhouse AM, Mets T, Brodiazhenko T, Abdullah M, Studer G, et al. Uncovering new families and folds in the natural protein universe. Nature. 2023;622:646–53.

39. Sunagawa S, Coelho LP, Chaffron S, Kultima JR, Labadie K, Salazar G, et al. Structure and function of the global ocean microbiome. Science. 2015;348:1261359.

40. Blondel VD, Guillaume J-L, Lambiotte R, Lefebvre E. Fast unfolding of communities in large networks. J Stat Mech Theory Exp. 2008;2008:P10008.

41. Vanni C, Schechter MS, Acinas SG, Barberán A, Buttigieg PL, Casamayor EO, et al. Unifying the known and unknown microbial coding sequence space. eLife. 2022;11.

42. Iyer LM, Leipe DD, Koonin EV, Aravind L. Evolutionary history and higher order classification of AAA+ ATPases. J Struct Biol. 2004;146:11–31.

43. Hedglin M, Kumar R, Benkovic SJ. Replication Clamps and Clamp Loaders. Cold Spring Harb Perspect Biol. 2013;5:a010165.

44. O’Donnell M, Onrust R, Dean FB, chen M, Hurwitz J. Homology in accessory proteins of replicative polymerases—E.coli to humans. Nucleic Acids Res. 1993;21:1–3.

45. Chia N, Cann I, Olsen GJ. Evolution of DNA Replication Protein Complexes in Eukaryotes and Archaea. PLOS ONE. 2010;5:e10866.

46. Yao NY, O’Donnell ME. Evolution of replication machines. Crit Rev Biochem Mol Biol. 2016;51:135–49.

47. Li H, O’Donnell M, Kelch B. Unexpected new insights into DNA clamp loaders. Bioessays. 2022;44:2200154.

48. Barre F-X, Søballe B, Michel B, Aroyo M, Robertson M, Sherratt D. Circles: The replication-recombination-chromosome segregation connection. Proc Natl Acad Sci. 2001;98:8189–95.

49. Romero H, Rösch TC, Hernández-Tamayo R, Lucena D, Ayora S, Alonso JC, et al. Single molecule tracking reveals functions for RarA at replication forks but also independently from replication during DNA repair in Bacillus subtilis. Sci Rep. 2019;9:1997.

50. Frickey T, Lupas AN. Phylogenetic analysis of AAA proteins. J Struct Biol. 2004;146:2– 10.

51. Lapointe F-J, Lopez P, Boucher Y, Koenig J, Bapteste E. Clanistics: a multi-level perspective for harvesting unrooted gene trees. Trends Microbiol. 2010;18:341–7.

52. Mirdita M, Schütze K, Moriwaki Y, Heo L, Ovchinnikov S, Steinegger M. ColabFold: making protein folding accessible to all. Nat Methods. 2022;19:679–82.

53. Jumper J, Evans R, Pritzel A, Green T, Figurnov M, Ronneberger O, et al. Highly accurate protein structure prediction with AlphaFold. Nature. 2021;596:583–9.

54. Cobbe N, Heck MMS. The Evolution of SMC Proteins: Phylogenetic Analysis and Structural Implications. Mol Biol Evol. 2004;21:332–47.

55. Rybenkov VV, Herrera V, Petrushenko ZM, Zhao H. MukBEF, a Chromosomal Organizer. J Mol Microbiol Biotechnol. 2015;24:371–83.

56. Petrushenko ZM, She W, Rybenkov VV. A new family of bacterial condensins. Mol Microbiol. 2011;81:881–96.

57. Kamada K, Barillà D. Combing Chromosomal DNA Mediated by the SMC Complex: Structure and Mechanisms. BioEssays. 2018;40:1700166.

58. Badel C, Bell SD. Chromosome architecture in an archaeal species naturally lacking structural maintenance of chromosomes proteins. Nat Microbiol. 2024;9:263–73.

59. Soppa J. Prokaryotic structural maintenance of chromosomes (SMC) proteins: distribution, phylogeny, and comparison with MukBs and additional prokaryotic and eukaryotic coiled-coil proteins. Gene. 2001;278:253–64.

60. Waldman VM, Stanage TH, Mims A, Norden IS, Oakley MG. Structural mapping of the coiled-coil domain of a bacterial condensin and comparative analyses across all domains of life suggest conserved features of SMC proteins. Proteins Struct Funct Bioinforma. 2015;83:1027–45.

61. Hirano T. The ABCs of SMC proteins: two-armed ATPases for chromosome condensation, cohesion, and repair. Genes Dev. 2002;16:399–414.

62. Gruber S, Arumugam P, Katou Y, Kuglitsch D, Helmhart W, Shirahige K, et al. Evidence that Loading of Cohesin Onto Chromosomes Involves Opening of Its SMC Hinge. Cell. 2006;127:523–37.

63. Connelly JC, Leach DRF. Tethering on the brink: the evolutionarily conserved Mre11– Rad50 complex. Trends Biochem Sci. 2002;27:410–8.

64. Kim E, Barth R, Dekker C. Looping the Genome with SMC Complexes. Annu Rev Biochem. 2023;92 Volume 92, 2023:15–41.

65. Lin Z, Kong H, Nei M, Ma H. Origins and evolution of the recA/RAD51 gene family: Evidence for ancient gene duplication and endosymbiotic gene transfer. Proc Natl Acad Sci. 2006;103:10328–33.

66. Thompson LH, Schild D. Homologous recombinational repair of DNA ensures mammalian chromosome stability. Mutat Res Mol Mech Mutagen. 2001;477:131–53.

67. Cox MM. The RecA protein as a recombinational repair system. Mol Microbiol. 1991;5:1295–9.

68. Seitz EM, Brockman JP, Sandler SJ, Clark AJ, Kowalczykowski SC. RadA protein is an archaeal RecA protein homolog that catalyzes DNA strand exchange. Genes Dev. 1998;12:1248–53.

69. Chintapalli SV, Bhardwaj G, Babu J, Hadjiyianni L, Hong Y, Todd GK, et al. Reevaluation of the evolutionary events within recA/RAD51 phylogeny. BMC Genomics. 2013;14:240.

70. Banerjee A, Neiner T, Tripp P, Albers S-V. Insights into subunit interactions in the Sulfolobus acidocaldarius archaellum cytoplasmic complex. FEBS J. 2013;280:6141–9.

71. Chaudhury P, Does C van der, Albers S-V. Characterization of the ATPase FlaI of the motor complex of the Pyrococcus furiosus archaellum and its interactions between the ATP-binding protein FlaH. PeerJ. 2018;6:e4984.

72. Castelle CJ, Brown CT, Anantharaman K, Probst AJ, Huang RH, Banfield JF. Biosynthetic capacity, metabolic variety and unusual biology in the CPR and DPANN radiations. Nat Rev Microbiol. 2018;16:629–45.

73. Schulz F, Roux S, Paez-Espino D, Jungbluth S, Walsh DA, Denef VJ, et al. Giant virus diversity and host interactions through global metagenomics. Nature. 2020;578:432–6.

74. Wu D, Wu M, Halpern A, Rusch DB, Yooseph S, Frazier M, et al. Stalking the Fourth Domain in Metagenomic Data: Searching for, Discovering, and Interpreting Novel, Deep Branches in Marker Gene Phylogenetic Trees. PLOS One. 2011;6.

75. Makarova KS, Krupovic M, Koonin EV. Evolution of replicative DNA polymerases in archaea and their contributions to the eukaryotic replication machinery. Front Microbiol. 2014;5.

76. Lennon CW, Stanger M, Banavali NK, Belfort M. Conditional Protein Splicing Switch in Hyperthermophiles through an Intein-Extein Partnership. mBio. 2018;9:10.1128/mbio.02304-17.

77. Sayers EW, Bolton EE, Brister JR, Canese K, Chan J, Comeau DC, et al. Database resources of the national center for biotechnology information. Nucleic Acids Res. 2022;50:D20–6.

78. Buchfink B, Reuter K, Drost H-G. Sensitive protein alignments at tree-of-life scale using DIAMOND. Nat Methods. 2021;18:366–8.

79. Hagberg AA, Schult DA, Swart PJ. Exploring Network Structure, Dynamics, and Function using NetworkX. In: Varoquaux G, Vaught T, Millman J, editors. Proceedings of the 7th Python in Science Conference. Pasadena, CA USA; 2008. p. 11–5.

80. Huerta-Cepas J, Szklarczyk D, Heller D, Hernández-Plaza A, Forslund SK, Cook H, et al. eggNOG 5.0: a hierarchical, functionally and phylogenetically annotated orthology resource based on 5090 organisms and 2502 viruses. Nucleic Acids Res. 2019;47:D309–14.

81. Spielman SJ, Wilke CO. Pyvolve: A Flexible Python Module for Simulating Sequences along Phylogenies. PLOS ONE. 2015;10:e0139047.

82. Shannon P, Markiel A, Ozier O, Baliga NS, Wang JT, Ramage D, et al. Cytoscape: a software environment for integrated models of biomolecular interaction networks. Genome Res. 2003;13:2498–504.

83. Li W, Godzik A. Cd-hit: a fast program for clustering and comparing large sets of protein or nucleotide sequences. Bioinforma Oxf Engl. 2006;22:1658–9.

84. Katoh K, Standley DM. MAFFT Multiple Sequence Alignment Software Version 7: Improvements in Performance and Usability. Mol Biol Evol. 2013;30:772–80.

85. Capella-Gutiérrez S, Silla-Martínez JM, Gabaldón T. trimAl: a tool for automated alignment trimming in large-scale phylogenetic analyses. Bioinformatics. 2009;25:1972–3.

86. Minh BQ, Schmidt HA, Chernomor O, Schrempf D, Woodhams MD, von Haeseler A, et al. IQ-TREE 2: New Models and Efficient Methods for Phylogenetic Inference in the Genomic Era. Mol Biol Evol. 2020;37:1530–4.

87. Hoang DT, Chernomor O, von Haeseler A, Minh BQ, Vinh LS. UFBoot2: Improving the Ultrafast Bootstrap Approximation. Mol Biol Evol. 2018;35:518–22.

88. Kalyaanamoorthy S, Minh BQ, Wong TKF, von Haeseler A, Jermiin LS. ModelFinder: fast model selection for accurate phylogenetic estimates. Nat Methods. 2017;14:587–9.

89. Zielezinski A, Vinga S, Almeida J, Karlowski WM. Alignment-free sequence comparison: benefits, applications, and tools. Genome Biol. 2017;18:1–17.

90. Ren J, Bai X, Lu YY, Tang K, Wang Y, Reinert G, et al. Alignment-Free Sequence Analysis and Applications. Annu Rev Biomed Data Sci. 2018;1:93–114.

91. Chan CX, Bernard G, Poirion O, Hogan JM, Ragan MA. Inferring phylogenies of evolving sequences without multiple sequence alignment. Sci Rep. 2014;4:6504.

92. Simonsen M, Mailund T, Pedersen CNS. Rapid Neighbour-Joining. In: Crandall KA, Lagergren J, editors. Algorithms in Bioinformatics. Berlin, Heidelberg: Springer; 2008. p. 113–22.

93. Letunic I, Bork P. Interactive Tree Of Life (iTOL) v5: an online tool for phylogenetic tree display and annotation. Nucleic Acids Res. 2021;49:W293–6.

94. Varadi M, Anyango S, Deshpande M, Nair S, Natassia C, Yordanova G, et al. AlphaFold Protein Structure Database: massively expanding the structural coverage of protein-sequence space with high-accuracy models. Nucleic Acids Res. 2022;50:D439–44.

95. The UniProt Consortium. UniProt: the Universal Protein Knowledgebase in 2023. Nucleic Acids Res. 2023;51:D523–31.

96. van Kempen M, Kim SS, Tumescheit C, Mirdita M, Lee J, Gilchrist CLM, et al. Fast and accurate protein structure search with Foldseek. Nat Biotechnol. 2023;:1–4.

