## Supplementary Material for "New groups of highly divergent proteins in families as old as cellular life with important biological functions in the ocean"

**Supplementary Text SI-1: Exploring the protein universe using iterative network searches unveiled distant homologs, including distant paralogs**

Sequences matched in earlier rounds of metagenome mining generally tended to have higher similarity to their closest *nr* relatives (one-sided Spearman rank test, correlation coefficient -0.2, p-value <$1.6x10^{-22}$) (Figure SI-8), highlighting the capacity of iterative alignment search protocols to identify distant homology relationships beyond the scope of direct BLAST searches. These earlier sequences also lay closer to seed sequences in sequence similarity networks, using alignment similarity as a network distance. In accordance with this, the network distance from environmental sequences to their seed counterparts was generally correlated with their divergence from their closest well-characterised published relative (one-sided Spearman test, correlation 0.22, p-value <$1.6x10^{-22}$). This observation stood both for the globality of environmental sequences and for a majority (47/53) of protein families when evaluated individually. Families that did not support this correlation exhibited a common pattern of environmental homolog retrieval, wherein an initial decline in retrieved sequences after each search step was followed by a substantial retrieval uptick, as opposed to an expected monotonous decline. Sequences responsible for this later retrieval uptick were generally more similar to their published counterparts than previously retrieved homologs, but connected only distantly to the seed sequences they were retrieved from.

Moreover, sequences from these 6 families were predominantly assigned to different OGs than previously retrieved homologs, suggesting that they may belong to other protein families than that initially used to query the environmental dataset, and showing that extant protein families do not always perfectly segregate in sequence homology within the protein space. Although these indirect connections may arise between bona fide unrelated families, spuriously showing some convergence in primary sequence, some may also be indicative of a shared evolutionary history.

One such case is for the SecD and SecF translocation proteins families. Both families are involved in a common complex facilitating protein secretion across the cytoplasmic membrane of archaea and bacteria [1]. While these two families had been separately retained by our clustering analysis of reference sequences as highly conserved and broadly distributed in accordance with the literature [2], our detection of distant homologs recovered their paralogy relationship, unveiling 11,864 sequences retrieved in common for both families out of 15,027 and 13,048 homologs in OM-RGC (in SecD and SecF respectively, after respectively 14 and 13 search iterations). Upon further analysis, these sequences clustered in two main categories. First-type sequences were closer to seed sequences from SecD than from SecF in the similarity networks, more central in the SecD network than in SecF, and retrieved by SecD sequences in fewer rounds of search; conversely, second-type sequences were closer to SecF by those same criteria. In line with these observations, eggNOG functional prediction [3] predominantly assigned first- and second-type sequences to OGs corresponding respectively to SecD (COG0342) and SecF (COG0341). Overall, nearly 80% of these sequences could be unambiguously mapped back to SecD or SecF by network distance, centrality, search iterations and eggNOG functional annotation. The intersecting retrieval of environmental homologs in the case of SecD and SecF proteins is thus most likely the result of their ancestral paralogous history [4], which was not reflected in the initial dataset but became apparent with the addition of environmental sequences, and assesses the efficiency of our iterative strategy to identify deep homology.


1. Pogliano JA, Beckwith J. SecD and SecF facilitate protein export in Escherichia coli. *EMBO J* 1994; **13**: 554–561.
2. Hand NJ, Klein R, Laskewitz A, Pohlschröder M. Archaeal and bacterial SecD and SecF homologs exhibit striking structural and functional conservation. *J Bacteriol* 2006; **188**: 1251–1259.

3. Huerta-Cepas J, Szklarczyk D, Heller D, Hernández-Plaza A, Forslund SK, Cook H, et al. eggNOG 5.0: a hierarchical, functionally and phylogenetically annotated orthology resource based on 5090 organisms and 2502 viruses. *Nucleic Acids Res* 2019; **47**: D309–D314.
4. Tseng T-T, Gratwick KS, Kollman J, Park D, Nies DH, Goffeau A, et al. The RND Permease Superfamily: An Ancient, Ubiquitous and Diverse Family that Includes Human Disease and Development Proteins. *J Mol Microbiol Biotechnol* 1999; **1**: 107–125.

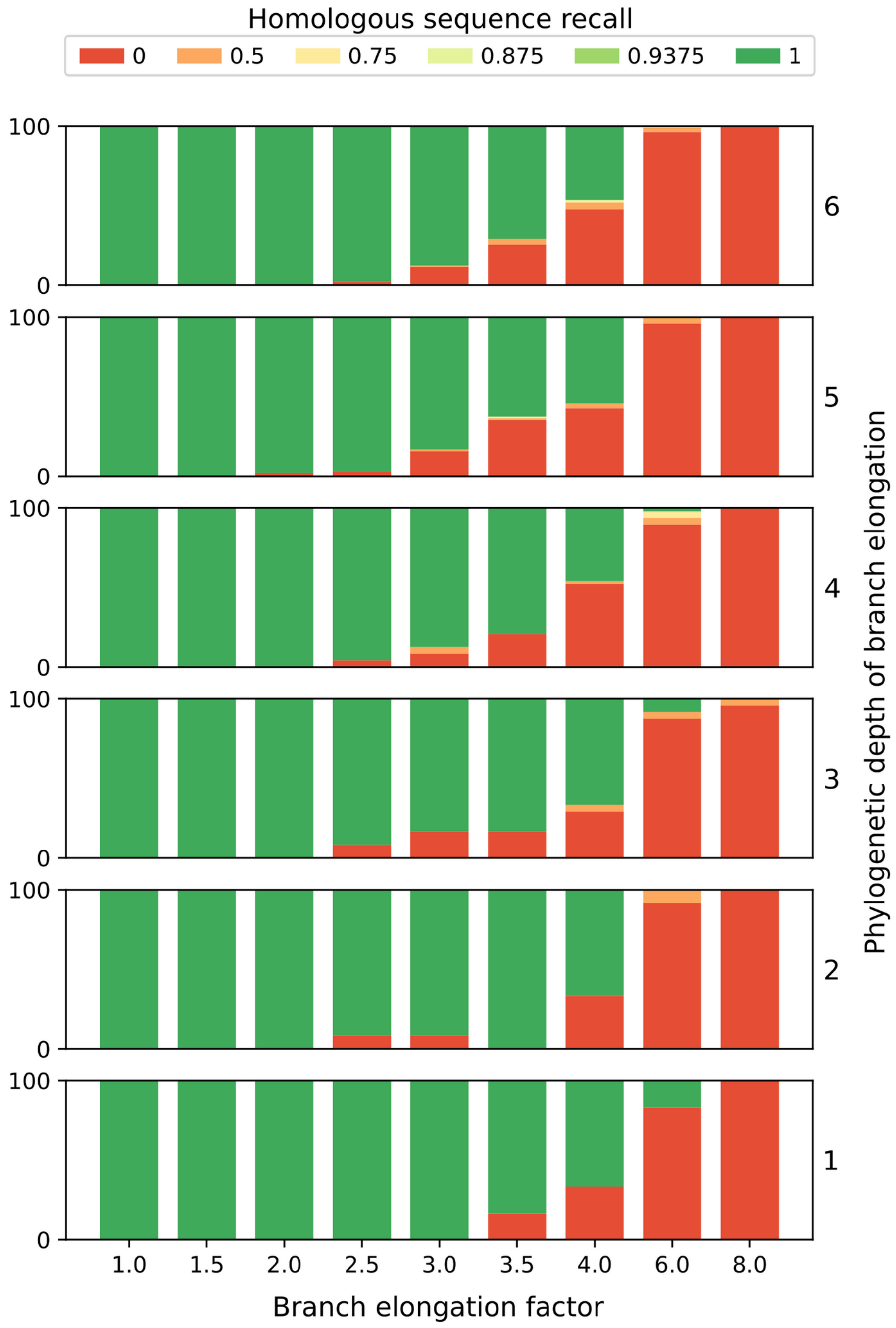


**Supplementary Figure SI-1: Distribution of the statistical recall value achieved by distant homology searches on simulated gene families.** Gene family simulations are grouped according to two simulation parameters: the branch elongation factor, quantifying the mutation rate increase between fast- and slow-evolving sequences, and the phylogenetic depth covered by the branch elongation (see Methods). Recall distributions are represented by stacked colour bars, and range from 0 (no divergent homolog retrieved) in red to 1 (all divergent homologs retrieved) in green.

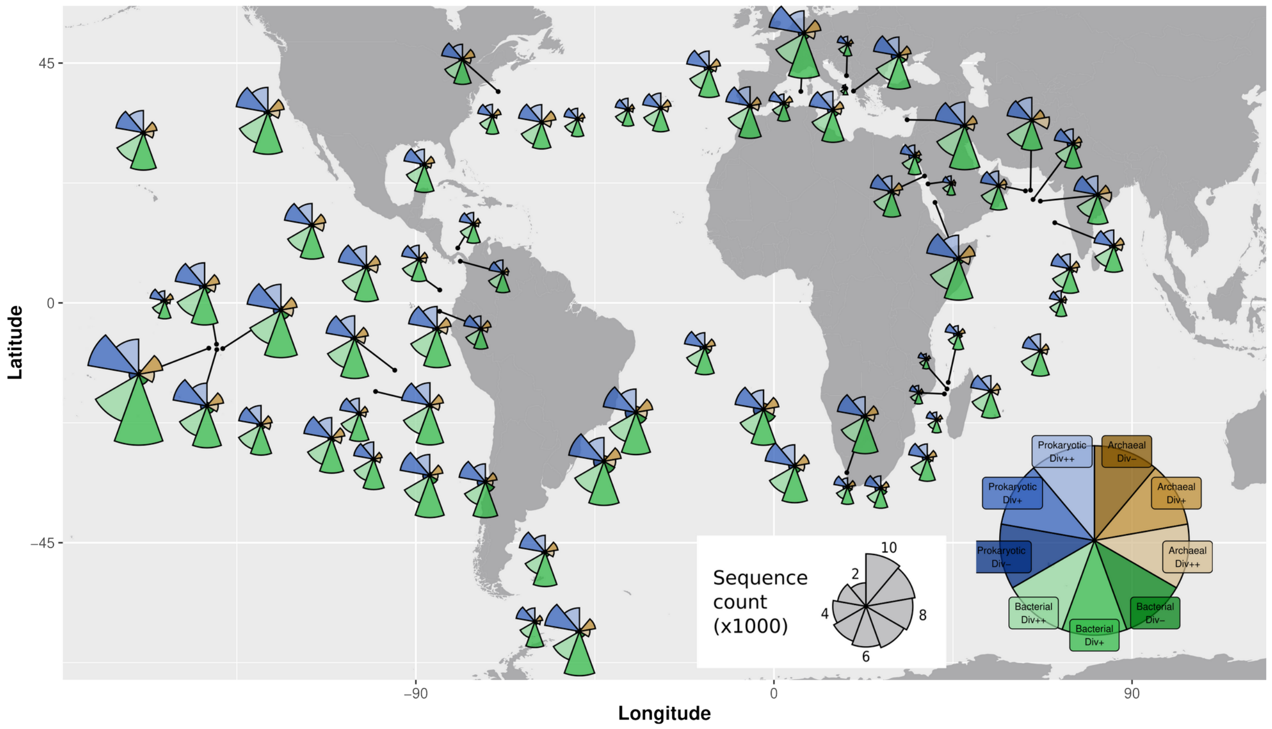


**Supplementary Figure SI-2: Geographical distribution of retrieved environmental homologs.** Environmental homologs were categorised as bacterial (green), archaeal (brown), or prokaryotic (blue), from highly divergent (Div++, <40% similarity with the best hit from cultured organism in the NCBI nr database) to weakly divergent (Div-, >90% similarity with the best hit from cultured organism in the NCBI nr database). The sizes of sector charts indicate the numbers of sequences falling within each category at a given sampling site.

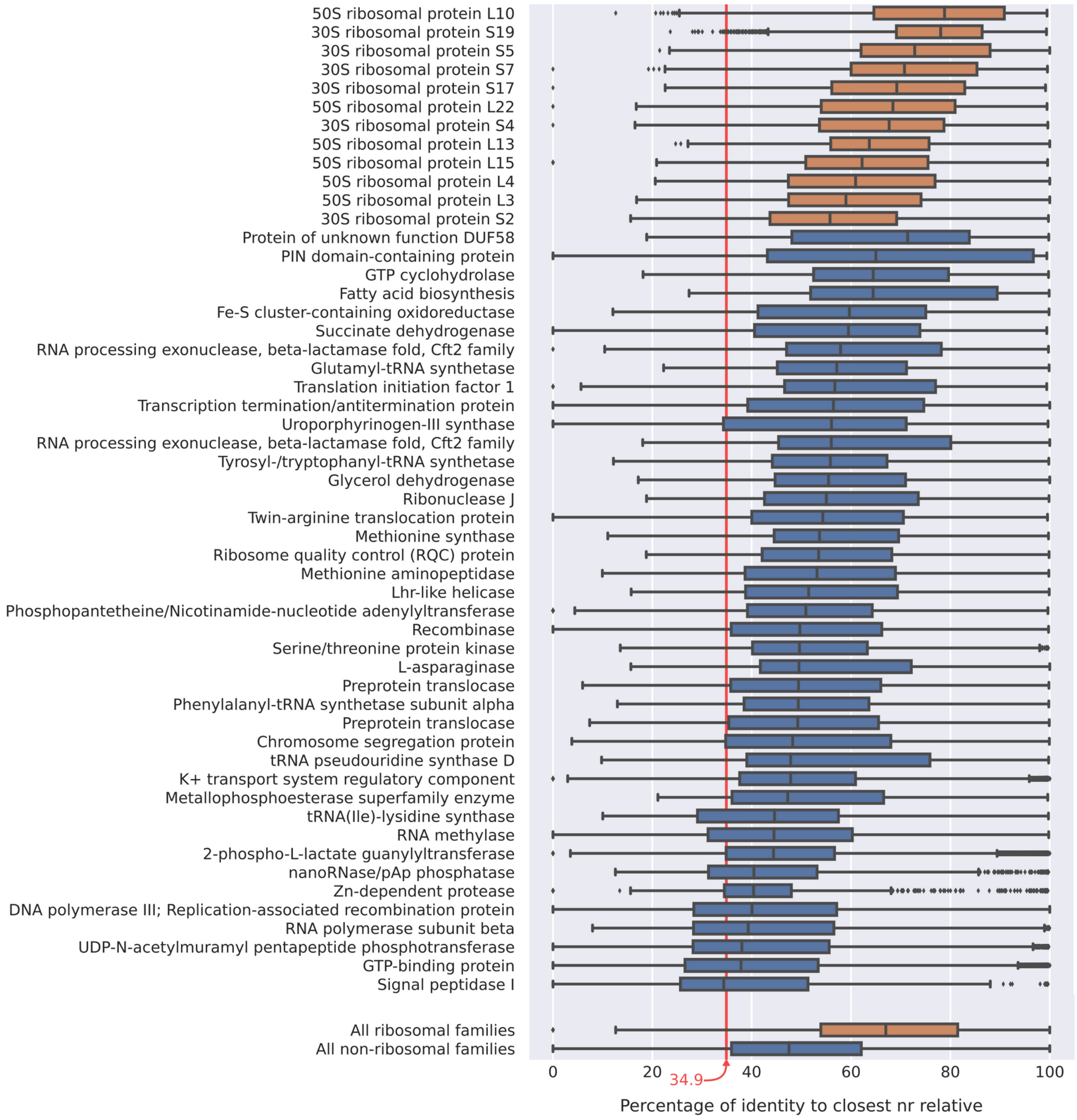


**Supplementary Figure SI-3: Distribution of the divergence of environmental homologs from the known diversity.** Boxplots show, for each of the 53 protein families, the distribution of sequence similarity between environmental homologs and their closest relative in the known cultured diversity (computed as the product of the amino-acid identity within the aligned region times the coverage of the alignment on the shortest sequence, ranging from 0% to 100%; environmental sequences with no known cultured relative are given a 0% score by default). Ribosomal families are represented by brown boxes, and blue boxes indicate non-ribosomal families. The two boxes on the bottom show the distribution of sequence similarity when all ribosomal families are pooled together against all non-ribosomal families. The red vertical line indicates the 34.9% identity threshold, corresponding to the average divergence observed between archaeal and bacterial sequences within seed protein families.

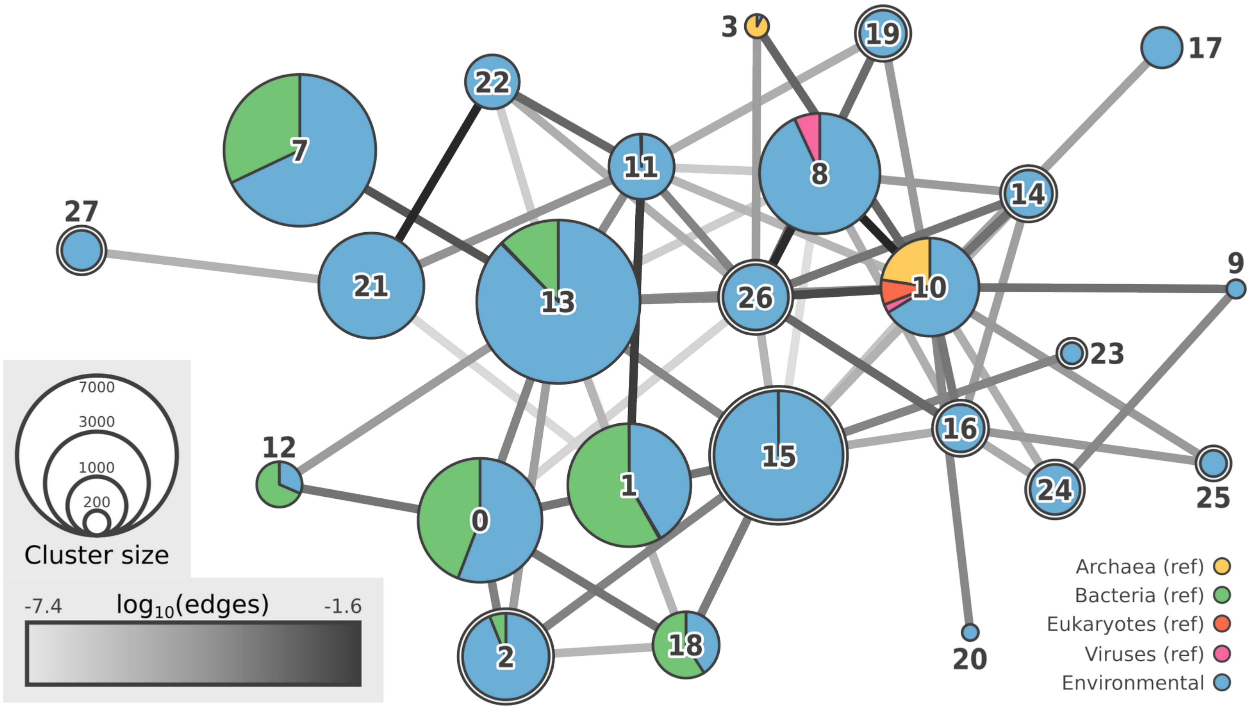


**Supplementary Figure SI-4: Meta-network of the clamp loader subunits sequences and their environmental homologs.** Nodes correspond to Louvain clusters within the sequence similarity network of clamp loader subunit proteins. Nodes are arbitrarily designated with numeric labels. Node areas indicate the number of represented sequences. The interior of each node consists of a pie chart indicating the composition of the cluster in archaeal, bacterial, eukaryotic, viral and environmental sequences. Nodes representing clusters of highly divergent environmental variants are highlighted with a double outline. Edges are coloured according to the proportion of edges between two clusters in the family SSN, relative to the maximum number of possible edges between these clusters (darker edges indicate stronger connections between clusters).

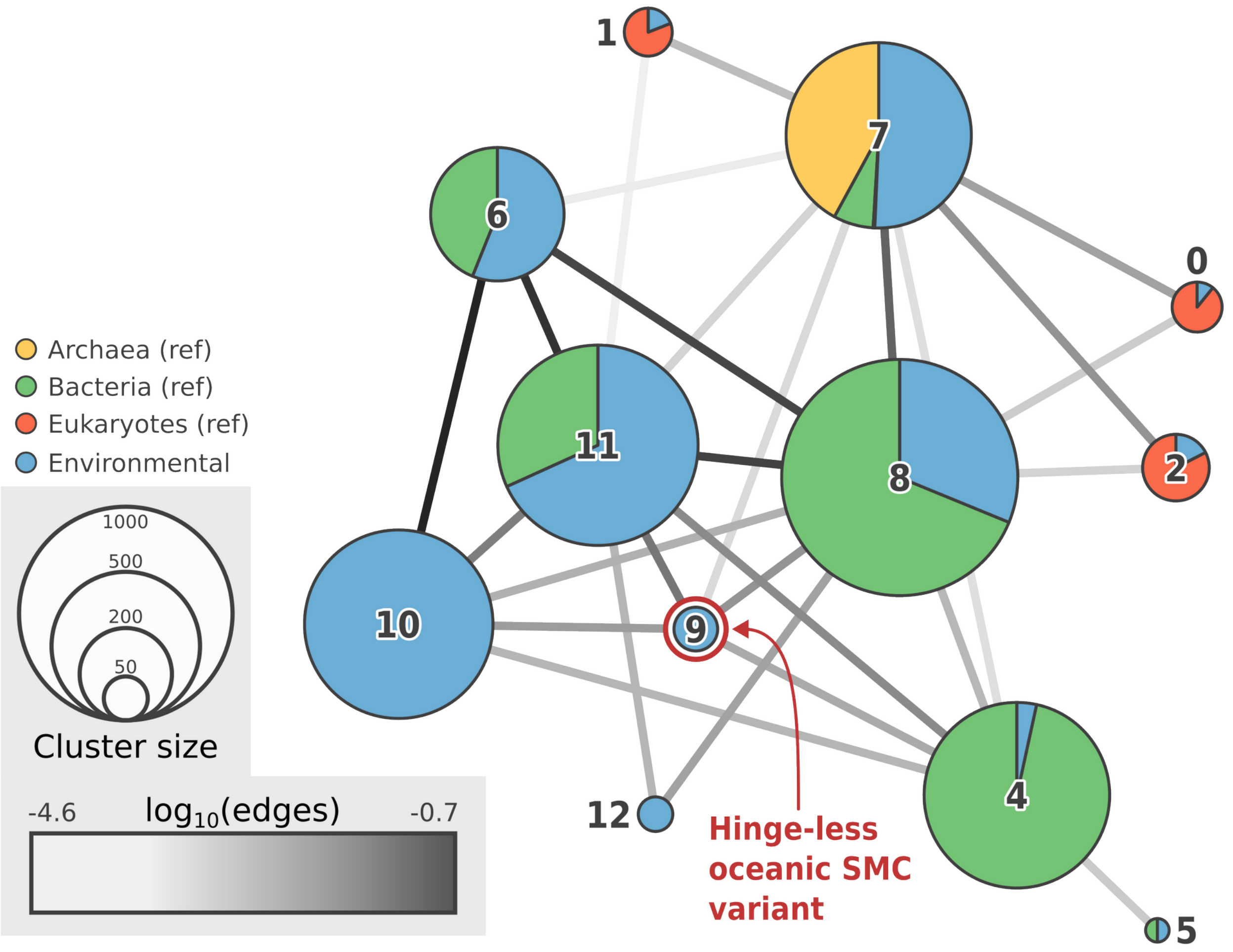


**Supplementary Figure SI-5: Meta-network of the SMC sequences and their environmental homologs.** Nodes correspond to Louvain clusters within the sequence similarity network of SMC proteins. Nodes are arbitrarily designated with numeric labels. Node areas indicate the number of represented sequences. The interior of each node consists of a pie chart indicating the composition of the cluster in archaeal, bacterial, eukaryotic and environmental sequences. Node 9, representing the only cluster of highly divergent environmental variants in this family, is highlighted in red. Edges are coloured according to the proportion of edges between two clusters in the family SSN, relative to the maximum number of possible edges between these clusters (darker edges indicate stronger connections between clusters).

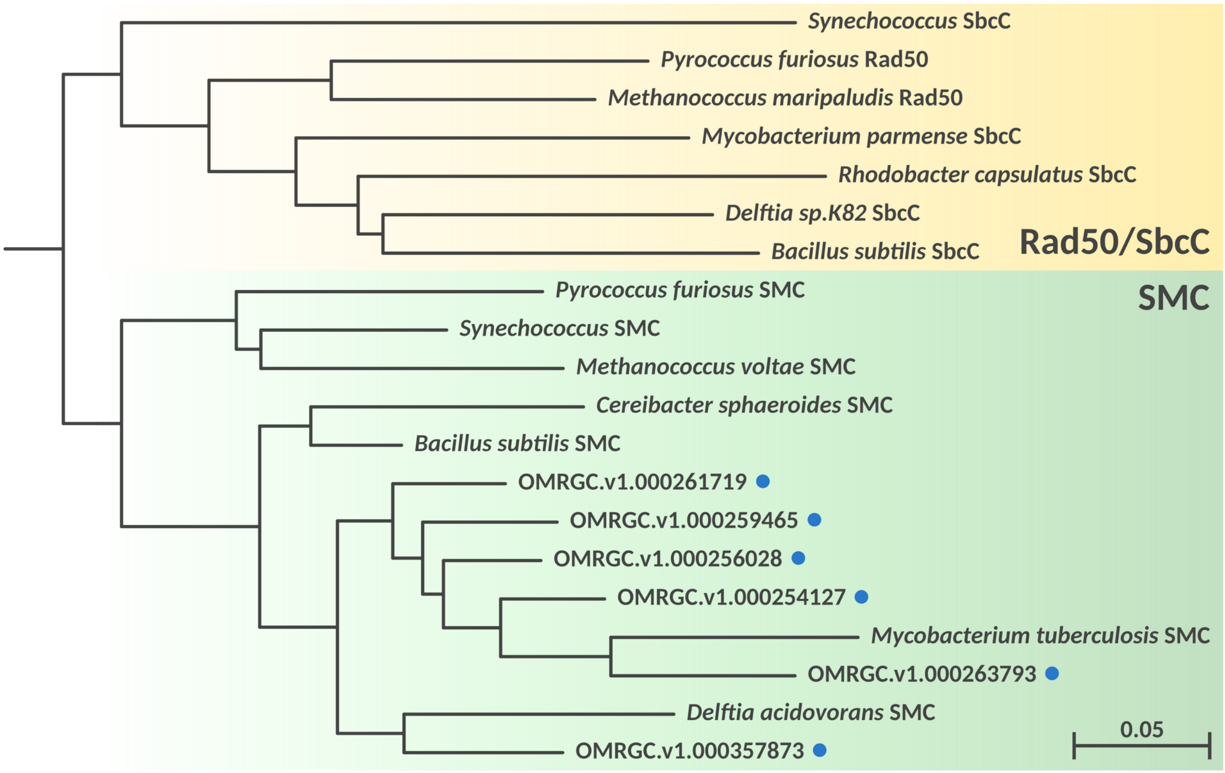


**Supplementary Figure SI-6: Dendrogram of tertiary structures of reference SMC proteins, environmental SMC homologs, and reference Rad50/SbcC proteins.** Protein structures were inferred with AlphaFold and compared (all against all) using Foldseek. The tree was rooted using midpoint-rooting, delimiting a SMC clade (green) and a Rad50/SbcC clade (yellow). OM-RGC SMC homologs are labelled by their database identifier and indicated by blue dots.

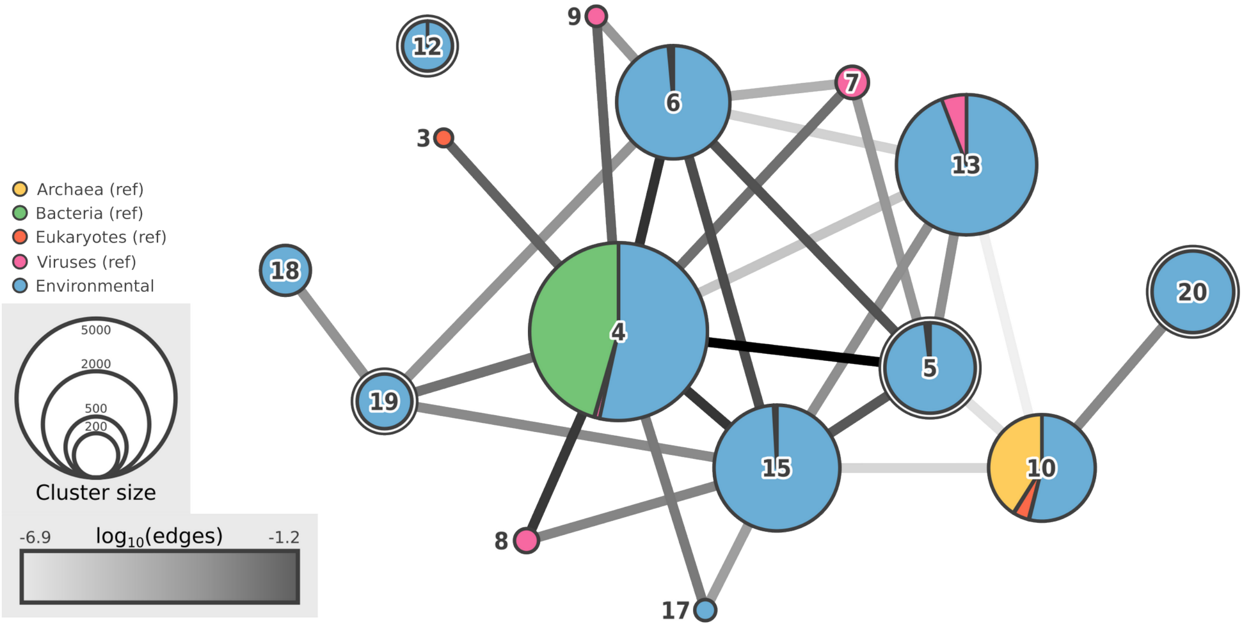

**Supplementary Figure SI-7: Meta-network of the RecA/RadA sequences and their environmental homologs.** Nodes correspond to Louvain clusters within the sequence similarity network of RecA/RadA proteins. Nodes are arbitrarily designated with numeric labels. Node areas indicate the number of represented sequences. The interior of each node consists of a pie chart indicating the composition of the cluster in archaeal, bacterial, eukaryotic, viral and environmental sequences. Nodes representing clusters of highly divergent environmental variants are highlighted with a double outline. Edges are coloured according to the proportion of edges between two clusters in the family SSN, relative to the maximum number of possible edges between these clusters (darker edges indicate stronger connections between clusters).

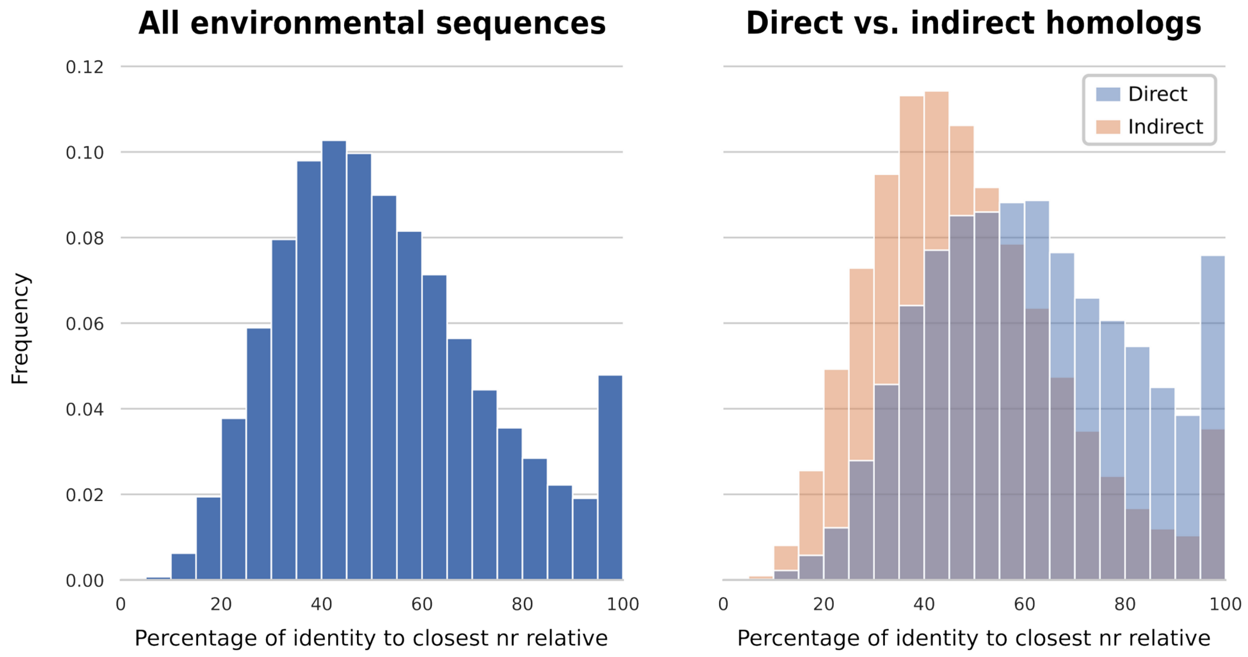


**Supplementary Figure SI-8: Distribution of sequence similarity to closest published homolog for environmental sequences.** Similarity is computed as amino-acid identity within the aligned region times coverage of the alignment on the shortest sequence, ranging from 0 to 100, and is indicated in abscissa. The proportion of environmental sequences with a given identity to their closest published relative is indicated in ordinate. **Left**: all environmental sequences. **Right**: sequences retrieved in the first round of search (blue) and in subsequent rounds (orange).

**Supplementary Table SI-1**: Public genomic data used to constitute the initial set of protein families. NCBI Assembly or Nucleotide IDs are indicated along with taxonomic information on the host lineage. This dataset was assembled from two prior separate sets of genome records, one with sequences linked to a Nucleotide identifier (sheet *Dataset_part1* in the Table) and one linking sequences to Assembly objects (sheet *Dataset_part2*), hence the two sheets in this Table.

**Supplementary Table SI-2**: Main functional annotations for 53 protein families used in the environmental homolog search. Families are specified by the predominant gene name(s) of sequences in the family. The size of the family before and after the iterative homolog gathering is indicated, along with the number of iterations required to gather all homologs.

**Supplementary Table SI-3**: Sequence identifiers, iteration step of retrieval, and SSN cluster label for seed sequences and environmental homologs of clamp loader subunits, SMC proteins, and recombinases. Seed sequences are identified in accordance with the seed dataset shared in Data Availability Statement and are designated as retrieved at step 0. Environmental sequences (retrieved at steps 1 and above) are specified by their OM-RGC identifiers.
